# Discovery of microsized RNA-derived RNAs and polypeptides in human cells

**DOI:** 10.1101/2023.04.24.538138

**Authors:** Xiaoqiang Huang, Haidong Wu, Weilong Zhong, Wenbo Li, Zhiyong Liu, Zhen Yang, Min Zhao, Xiaonan Xi, Bo Cao, Yong Pu, Xiangxu Kong, Huan Zhao, Ronghua Zhang, Keguan Lai, Xinze Lv, Yue Lv, Jingyu Bao, Meimei Wang, Yanan Xiong, Xiubin Liang, Xiaokang Wu, Zhuying Wei, Youcai Xiong, Juan Zhong, Jifeng Zhang, Gilbert S. Omenn, Yuanjing Hu, Jie Xu, Guangling Zhang, Y. Eugene Chen, Shuang Chen

## Abstract

MicroRNAs (miRNAs) are short noncoding RNAs that can regulate gene expression through the binding of their 5’-ends to mRNAs. However, the biological effects of miRNA binding via their 3’-ends remain unclear. Here, we discover that the exact reverse pairing of the 3’-ends of miRNAs or miRNA-like RNAs, collectively termed microsized RNAs (msRNAs), with template RNAs can initiate the production of msRNA-derived RNAs (msdRs), which can subsequently be translated into polypeptides (msdPs). Using 2,632 human msRNAs from miRBase, 11,121 msdRs and 1,239 msdPs were predicted based on a 15-nucleotide pairing threshold, and the presence of representative msdRs and msdPs was confirmed in human cells. Of clinical relevance, msdP0188 is highly expressed in human lung and breast cancers, and its corresponding msRNAs and msdRs represent novel anti-cancer targets. Notably, inhibiting telomerase reverse transcriptase, a putative RNA-dependent RNA polymerase identified by bioinformatic screening, led to reduced levels of msdP0188 in human cancer cells. Our findings reveal a novel “msRNA → msdR → msdP” axis, expanding the classical central dogma and predicting many previously unrecognized RNAs and polypeptides with potential biological functions in human cells and other systems.

## Introduction

MicroRNAs (miRNAs) are short noncoding RNAs that play a crucial role in post-transcriptional gene regulation, influencing various biological processes ^1–6^. It is well established that miRNAs exert their regulatory functions by binding their 5’-ends to complementary sequences on target mRNAs ^7–9^. This naturally raises an intriguing question: could the pairing of a miRNA’s 3’-end with a complementary RNA induce biological effects? To our knowledge, this possibility remains largely unexplored.

Inspired by the RNA replication mechanism of RNA-dependent RNA polymerase (RdRP) in viruses ^10,11^, we hypothesize that, by pairing its 3’-end with a template RNA, a miRNA or miRNA-like RNA, collectively termed microsized RNA (msRNA), can act as a primer to initiate the synthesis of novel RNAs (<u>ms</u>RNA-<u>d</u>erived <u>R</u>NAs, msdRs) that are unlikely to arise through conventional transcription. We further postulate that msdRs may encode polypeptides (msdPs).

Here, we bioinformatically predicted msdRs and msdPs followed by experimental validation of the presence of selected msdRs and msdPs in human cells. We show that msdP0188, a representative msdP, is highly enriched in lung and breast cancer tissues and that its corresponding msRNAs and msdRs represent novel anti-cancer targets. Additionally, we find that telomerase reverse transcriptase (TERT) may function as an RdRP in producing msdRs and msdPs. This unconventional “msRNA → msdR → msdP” pathway not only extends the classical central dogma but also potentially reveals many unrecognized RNAs and polypeptides with potential biological and pathological functions in human cells and other systems.

## Results

### Prediction of msdRs and msdPs

The overall workflow for predicting msdRs and msdPs is shown in **Fig. 1A**. Briefly, the 3’-end sequence of a msRNA is searched against the NCBI RefSeq RNA database ^12^ using BLASTN ^13^ to find complementary RNAs; the msRNA sequence defines the 5’-end of a putative msdR, which is extended to the 5’-end of the template RNA. Potential internal ribosome entry sites (IRESs) are predicted by aligning the msdR with validated IRES sequences ^14^, assuming functionality if coverage and aligned identity thresholds are met (Methods). An msdP is predicted from any start codon located within 100 nucleotides (nt) downstream of a putative IRES, considering all combinations of multiple IRESs and start codons within each msdR.

**Fig. 1.**
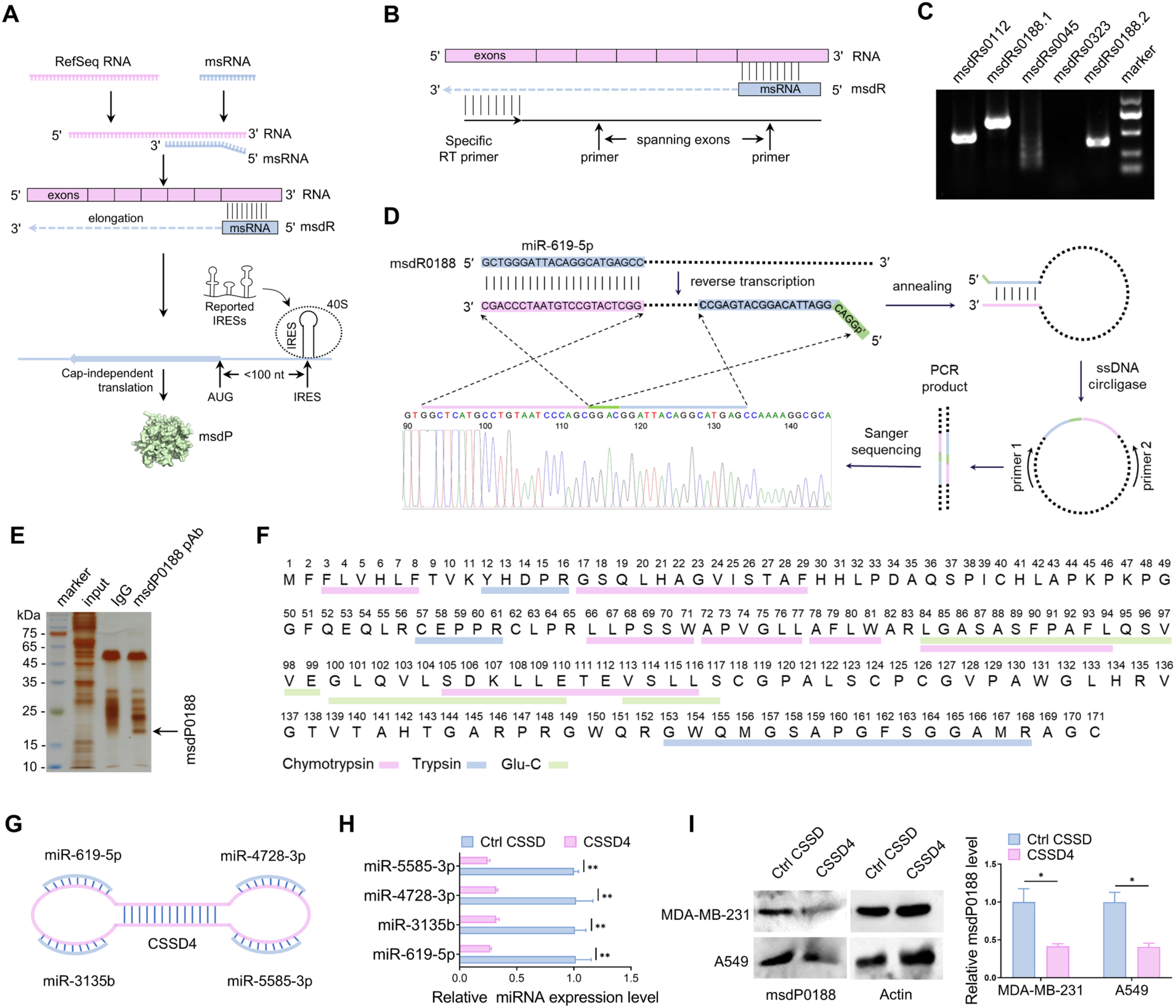
Prediction and validation of microsized RNA (msRNA)-derived RNAs (msdRs) and proteins (msdPs). **(A)** Bioinformatic pipeline for predicting msdRs and msdPs. (**B**) Schematic representation of the design of specific reverse transcription (RT) primers and exon-spanning PCR primers for msdRs. (**C**) Agarose gel electrophoresis analysis of msdR fragments amplified by specific exon-spanning primers. (**D**) Verification of the 5’-end sequence of msdRs0188 using ssDNA circligase-dependent 5’ RACE. (**E**) Silver staining of proteins separated by SDS-PAGE and captured using the anti-msdP0188 polyclonal antibody (pAb). The msdP0188 band is indicated. (**F**) Confirmation of msdP0188 through mass spectrometry analysis following digestion with trypsin, chymotrypsin, and Glu-C. (**G**) Schematic illustration of CSSD4, which simultaneously adsorbs four msRNAs (*miR-619-5p*, *miR-5585-3p*, *miR-4728-3p*, and *miR-3135b*) bound to *AP1M1* mRNA with a pairing length of ≥12 nt. (**H**) Expression levels of *miR-619-5p*, *miR-5585-3p*, *miR-4728-3p*, and *miR-3135b* in A549 cells after CSSD transfection. (**I**) Expression levels of msdP0188 in A549 and MDA-MB-231 cells after CSSD transfection. Error bars represent the standard error of the mean (s.e.m.) from n = 3 independent biological replicates. \**P* < 0.05; \*\**P* < 0.01 (Student’s *t*-test).

Using 2,632 human miRNAs from miRBase ^15^ as example msRNAs—given their short length and RNA-binding capacity—we applied a 15-nt pairing threshold and obtained 11,121 msdRs and 1,239 msdPs, excluding msdPs under 10 amino acids (**Table S1; Table S2; Data S1**). Additional predictions were performed with thresholds of 10–14 nt (**Table S2; Data S2; Data S3**).

To estimate the potential false discovery rate (FDR) for msdR prediction, we compared msdR predictions from the miRBase miRNA set (predicted positives, PP) with those from nine shuffled msRNA control sets (false positives, FP) using a shuffling method adopted from previous studies ^16,17^ (**Methods**; **Fig. S1**). For the 15-nt match, PP and FP were 11,121 and 1,695, respectively, yielding an FDR of ∼15%, confirming a stronger signal in miRBase miRNAs (**Table S3**). As expected, FDRs increased for shorter matches. Simulations using random msRNAs showed even lower FDRs (∼11%) (**Table S3**). Details on simulated msRNAs and msdR predictions are provided (**Data S4; Data S5**).

A systematic naming scheme was developed to assign identifiers to msdRs and msdPs, incorporating the msRNA name, template RNA GenInfo Identifier, gene symbol, and msRNA 3’-end matching position (Methods; **Data S2; Data S3**). For clarity, the 1,239 msdPs were also labeled sequentially (msdP0001– msdP1239) based on the ASCII order of their sequences. Notably, msRNAs, msdRs, and msdPs may not follow a one-to-one mapping: one msdR may produce multiple msdPs (**Fig. S2A**); distinct msRNAs targeting the same RNA may yield different msdRs but identical msdPs (**Fig. S2B**); and one msRNA may bind multiple RNAs or isoforms, producing varied msdR/msdP outputs (**Fig. S2C, D**).

Among the 1,239 msdPs, 46 matched sequences in the NCBI nr database ^12^ with high identity and coverage based on BLASTP ^13^ search, including six with 100% identity (Methods; **Data S6; Data S7**). The remaining 1,193 msdPs (96%) lacked intense matches and thus may represent novel polypeptides in human cells (**Data S8**).

### Presence of msdRs and msdPs in human cells

We then compared the 1,193 msdPs against mass spectrometry (MS) data from four tumor cell lines using Sequest-HT ^18^ (Methods; **Fig. S3**), identifying 941 msdPs (79%) with intense matches based on coverage, peptide-spectrum matches (PSMs), and Sequest-HT scores (**Data S9**). In parallel, PepQuery2 ^19^ analysis against over a billion MS/MS spectra in PepQueryDB identified high-confidence PSMs for 1,095 peptides from 555 msdPs (**Fig. S3; Data S10**), with 459 msdPs overlapping between two analyses.

To assess msdR expression, we selected four msdPs of varying lengths for further study in A549 cells (**Table S4; Table S5**). Primers were designed to amplify regions unique to the complementary DNA (cDNA) sequences of each putative msdRs, absent from genomic DNA (**Fig. 1B**). For msdRs0188, two fragments were amplified for dual verification; for the others, a single fragment was targeted (**Fig. 1C; Table S6**). Reverse transcription (RT)-PCR and Sanger sequencing confirmed the presence of msdRs0112 and msdRs0188 fragments but not msdRs0045 and msdRs0323 (**Fig. 1C; Data S11**).

To verify that msdRs arise from msRNA priming, we performed 5’ rapid amplification of cDNA ends (5’ RACE) to determine whether the 5’-end of msdRs0188 aligns precisely with its predicted msRNA source, *hsa-miR-619-5p* (Methods; **Fig. 1D**). A sequeence-specific primer was used to synthesize a 585-nt cDNA, which included terminal overhangs to enhance circularization efficiency. Following circularization and PCR amplification, Sanger sequencing of the junction site confirmed that the 5’-end of msdRs0188 matched the *hsa-miR-619-5p* sequence, supporting its derivation from this msRNA (**Fig. 1D; Data S11**).

To detect msdP expression in human tissues, peptide immunogens specific to msdPs were designed to generate rabbit polyclonal antibodies (pAbs) (**Table S4**), validated for sensitivity and specificity (Methods; figs. S4-S6). Immunohistochemistry (IHC) using these pAbs was employed to detect the expression of msdP0112 and msdP0188 in human cancer tissues and normal tissues. The anti-msdP0188 pAb led to intense, positive staining in lung and breast cancer tissues collected from various patients, and the anti-msdP0112 pAb resulted in robust positive staining in lung and colon cancer tissues (**Fig. 2; Fig. S7; Fig. S8**). These data suggest that both msdRs and msdPs are endogenously expressed in human tumor cells and may play a role in cancer biology.

**Fig. 2.**
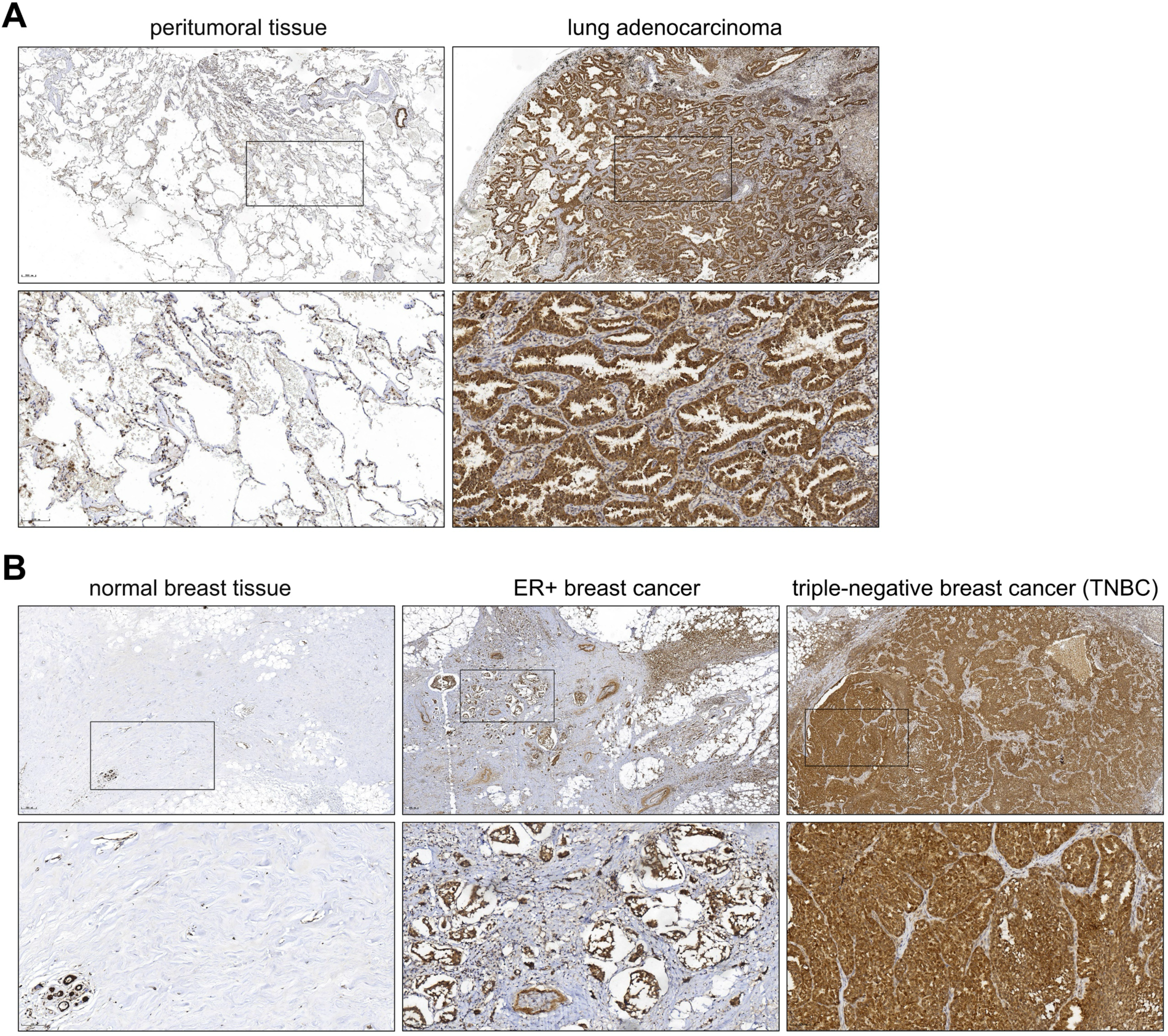
Elevated expression of msdP0188 in human lung and breast cancers. (**A**) Representative immunohistochemical staining of msdP0188 in normal lung tissues (n = 9) and lung cancer tissues (n = 15) using the anti-msdP0188 polyclonal antibody (1:200). Images are shown at 5× (upper) and 20× (lower) magnification. (**B**) Representative immunohistochemical staining of msdP0188 in normal breast tissues (n = 8), estrogen receptor-positive (ER+) breast cancer tissues (n = 10), and triple-negative breast cancer (TNBC) tissues (n = 12) using the anti-msdP0188 polyclonal antibody (1:200). Images are shown at 5× (upper) and 20× (lower) magnification.

### Bona fide confirmation of msdP0188 in human cells

A549 cell lysates were analyzed by SDS-PAGE and silver staining, and a band corresponding to ∼18 kDa, consistent with msdP0188, was identified (**Fig. 1E**). The band was excised, digested with three proteases, and subjected to MS/MS. This analysis identified 13 peptides spanning 55% of msdP0188’s sequence (94/171 amino acids) (**Fig. 1F; Fig. S9; Data S12**). BLASTP analysis of four extended peptides found no high-similarity hits in the NCBI nr database (**Table S7**), indicating these peptides likely represent new entities.

We next tested whether knockdown of the msRNAs predicted to give rise to msdP0188 would reduce its expression. Using the stringent 15-nt matching threshold, *hsa-miR-619-5p* was identified as the primary msRNA binding to mRNA *AP1M1* variants 1 or 2. Relaxing the threshold to 12 nt identified three additional msRNAs: *hsa-miR-3135b*, *hsa-miR-4728-3p*, and *hsa-miR-5585-3p* (**Fig. S10**).

To simultaneously target all four msRNAs, we designed a circular single-stranded DNA (CSSD4), which functions as a molecular sponge ^20,21^ (**Fig. 1G; Fig. S11**). A control CSSD containing scrambled sequences was used for comparison. CSSD4 specificity was confirmed by showing no effect on unrelated msRNAs (*hsa-miR-21-5p*, *hsa-miR-15a-5p*, and *hsa-miR-34a-5p*) (**Fig. S12**). In contrast, CSSD4 considerably reduced all four target msRNAs in A549 cells (**Fig. 1H**), leading to decreased msdP0188 protein levels (**Fig. 1I**).

To determine if direct targeting of msdRs would similarly impact msdP0188, we generated stable MDA-MB-231 and A549 cell lines expressing short hairpin RNAs (shRNAs) against msdR sequences. Western blot analysis showed that shRNA-2 was the most effective, leading to a substantial reduction of msdP0188 levels in both cell lines (**Fig. S13A,B**).

Together, these results confirm that msdP0188 is expressed in human cells and is produced through a defined “msRNA → msdR → msdP” pathway.

### Targeting msdP0188 in lung and breast cancer cells in vitro and in vivo

We hypothesized that specific msdPs and their corresponding msRNAs and msdRs may serve as novel biomarkers or therapeutic targets in human cancers. To test this, we investigated whether msdP0188, its originating msRNAs, and its encoding msdRs could be viable targets for cancer intervention.

Transwell assays and wound healing assays indicated that knockdown of msdP0188-associated msdRNAs via CSSD4 effectively inhibited the invasive and migratory capabilities of MDA-MB-231 and A549 cells compared to the control CSSD groups (**Fig. 3A,B**). Scanning electron microscopy (SEM) imaging further revealed morphological changes indicative of altered cellular architecture following CSSD4 treatment (**Fig. 3C**). Similar inhibitory effects on invasion, migration, and cell morphology were observed upon knockdown of msdP0188-encoding msdRs using shRNA (**Fig. S13C-E**).

**Fig. 3.**
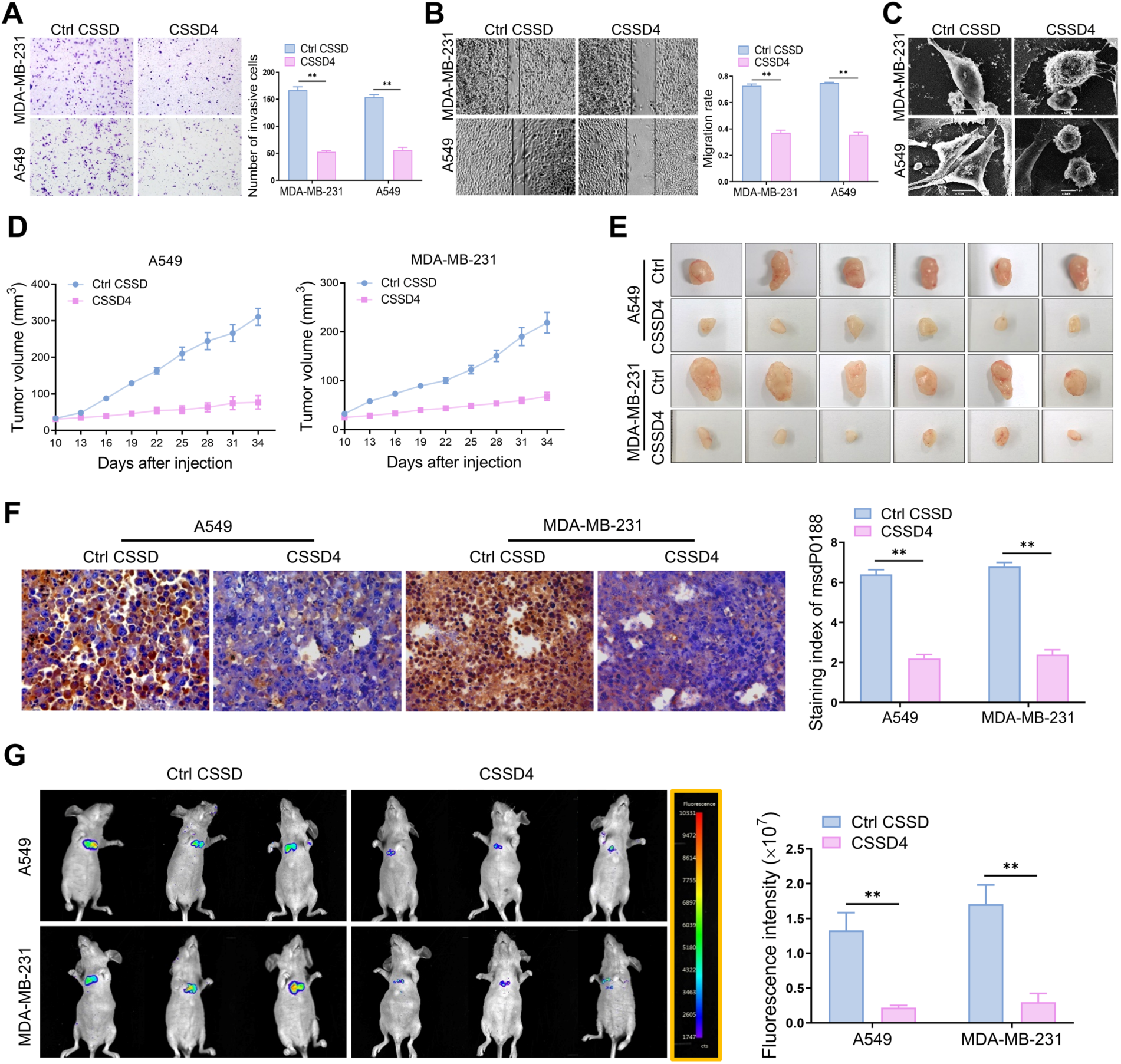
CSSD4 suppresses lung and breast cancer cell progression in vitro and in vivo. (**A**) Effect of CSSD on A549 and MDA-MB-231 cell invasion. (**B**) Effect of CSSD on A549 and MDA-MB-231 cell migration. (**C**) Scanning electron microscopy analysis of cell phenotype changes following CSSD transfection. (**D**) Solid tumors formed by A549 and MDA-MB-231 cells in nude mice following CSSD treatment. (**E**) Tumor volume measurements corresponding to Fig. 3D. (**F**) Immunohistochemical staining intensity of msdP0188 in solid tumors. (**G**) Lung metastasis of A549 and MDA-MB-231 cells following CSSD treatment. Error bars represent s.e.m. from n = 6 independent biological replicates in Fig. 3D; n = 3 in all other cases. \**P* < 0.05; \*\**P* < 0.01 (Student’s *t*-test).

Conversely, overexpression of msdP0188 in A549 cells promoted migratory and invasive ability relative to vector-transfected controls (**Fig. S14A-C**), accompanied by noticeable morphological alterations observed via SEM (**Fig. S14D**).

We extended these findings in vivo using two mouse models: a subcutaneous solid tumor xenograft model and a tail vein injection metastasis model. In the xenograft model, 24 nude mice were randomized into four groups and inoculated subcutaneously with A549 or MDA-MB-231 cells. Once tumors reached ∼4 mm in diameter, mice were intratumorally injected with CSSD4 or control CSSD every seven days for a total of three doses. Tumor sizes were measured every three days, and tumors in CSSD4-treated mice were noticeably smaller than those in the control groups for both cancer types (**Fig. 3D,E**). IHC analysis of excised tumors confirmed reduced msdP0188 expression in CSSD4-treated tissues (**Fig. 3F**).

In the metastasis model, 12 nude mice were randomly divided into four groups and injected intravenously with luciferase-expressing A549 or MDA-MB-231 cells. CSSD4 or control CSSD was administered intravenously on days 10, 17, and 24 post-injection. Bioluminescence imaging revealed a markedreduction in lung metastases in CSSD4-treated mice compared to controls (**Fig. 3G**), demonstrating suppressed metastatic potential following msdP0188-targeting intervention.

Collectively, these findings demonstrate that msdP0188 promotes the invasion and migration of lung and breast cancer cells. Both in vitro and in vivo evidence support that targeting msdP0188—either via its originating msRNAs or its encoding msdRs—effectively suppresses tumor progression. These results highlight msdP0188 as a promising therapeutic target for lung and breast cancers.

### TERT as a putative RdRP involved in msdR/msdP biogenesis

The detection of msdRs in human cells implies the presence of RdRP-like activity, traditionally thought to be absent in higher eukaryotes. We hypothesized that human proteins with features conserved among known RdRPs—either from eukaryotes, viruses, or DNA-dependent RNA polymerases (DdRPs)—might fulfill this role. To explore this, we conducted a comprehensive bioinformatic screening for candidate human RdRPs (**Fig. 4A**).

**Fig. 4.**
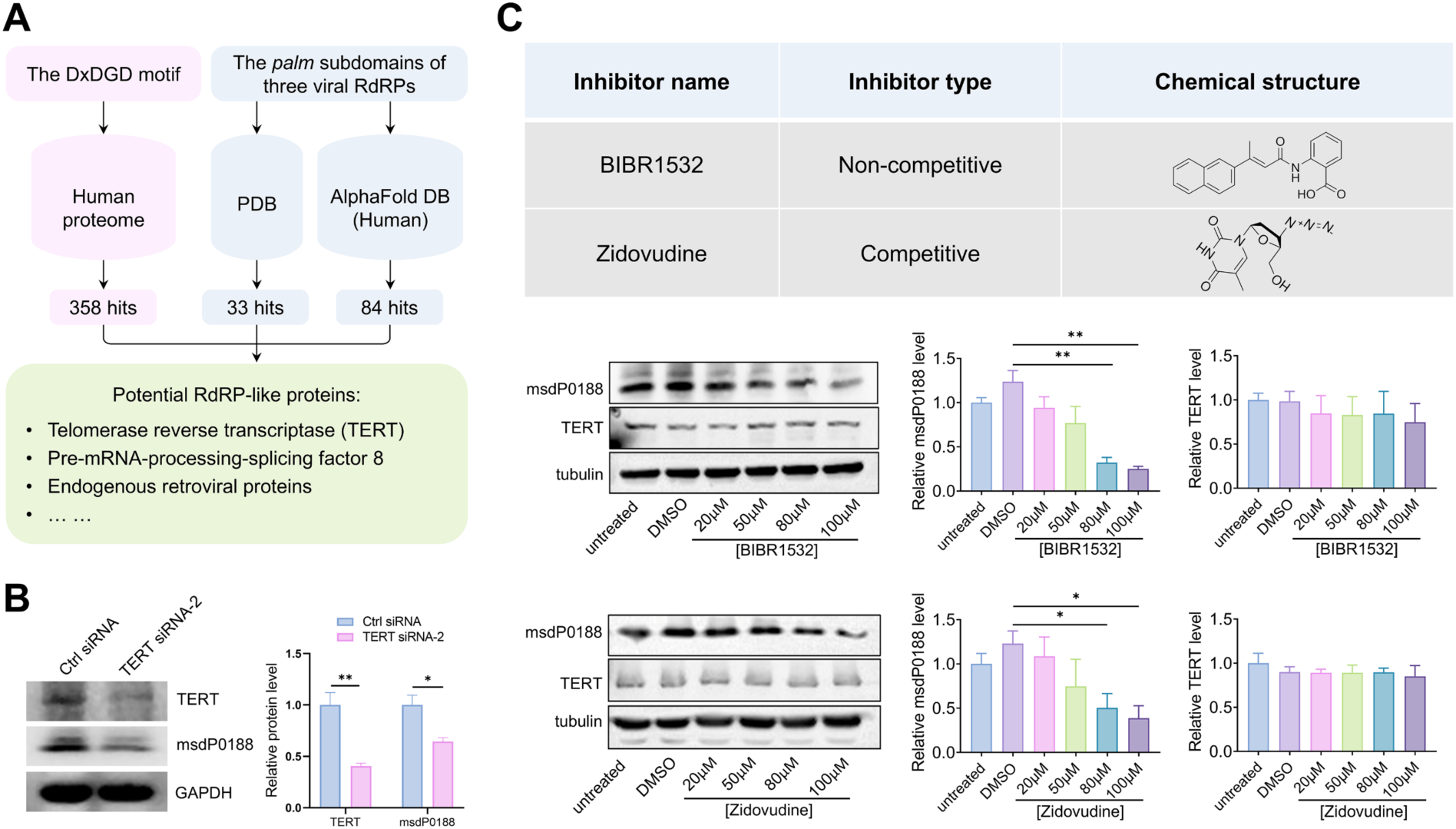
Human telomerase reverse transcriptase (TERT) is implicated in msdR generation. (**A**) Bioinformatic prediction of RdRP-like proteins involved in msdR production in human cells. (**B**) Relative expression levels of TERT and msdP0188 following TERT knockdown using siRNA-2 and control siRNA. (**C**) Relative expression levels of msdP0188 and TERT in A549 cells following treatment with TERT inhibitors BIBR1532 and zidovudine at different concentrations. Untreated and DMSO-treated A549 cells were used as controls. Error bars represent s.e.m. from n = 3 independent biological replicates. \**P* < 0.05; \*\**P* < 0.01 (Student’s *t*-test in Fig. 4B; one-way ANOVA with Tukey’s post-hoc test in Fig. 4C).

Eukaryotic RdRPs and the β’ subunit of DdRPs typically harbor a conserved catalytic D*x*DGD motif, where *x* denotes a bulky residue ^22^. Using this motif as a query and applying a length threshold of more than 500 amino acids, we identified 358 human proteins containing the D*x*DGD motif, many of which are known DdRPs or related subunits (**Data S13**).

In contrast, viral RdRPs generally lack the D*x*DGD motif but share a conserved catalytic architecture containing three subdomains: *fingers*, *palm*, and *thumb*, with the *palm* subdomain being structurally conserved. We assessed the structural similarity of the *palm* subdomains from three viruses (SARS-CoV-2, human picobirnavirus strain Hy005102, and *Thosea asigna* virus) using TM-align ^23^. All pairwise comparisons yielded TM-scores > 0.5, confirming a conserved structural fold across these viral RdRPs (**Fig. S15**). This observation supported the feasibility of identifying structurally analogous domains in the human proteome.

We searched for human proteins with structural similarity to the *palm* subdomains within the Protein Data Bank (PDB) ^24^ and the AlphaFold Protein Structure Database (AFDB) ^25,26^, applying a TM-score cutoff >0.5. This search yielded 33 candidate proteins from PDB and 84 from AFDB (**Data S14; Data S15**).

Among the top candidates was telomerase reverse transcriptase (TERT), a well-characterized reverse transcriptase that has previously been shown to possess RdRP activity ^27–30^. In addition to TERT, we found nine endogenous retroviral proteins that exhibit strong structural homology to the viral *palm* domains at the fold level (**Data S14; Data S15**). Given the elevated telomerase activity in many cancer cells ^31,32^ and the high levels of msdP0188 observed in lung and breast cancer cells, we hypothesized that TERT may participate in the biogenesis of msdRs and msdPs.

Initial attempt to knock out TERT in A549 cells using CRISPR/Cas9 resulted in extensive cell death, precluding further analysis. Therefore, we used small interfering RNAs (siRNAs) to knock down TERT transiently. Among the five siRNAs tested, siRNA-2 exhibited the most effective knockdown (**Fig. S16**). Notably, TERT knockdown using siRNA-2 led to an observable reduction in both TERT protein levels and msdP0188 expression (**Fig. 4B**), implicating TERT in msdP0188 biogenesis.

A549 cells were also treated with two established TERT inhibitors: BIBR1532 (a noncompetitive inhibitor) and Zidovudine (a competitive nucleoside analog). At high concentrations, both inhibitors substantially reduced msdP0188 protein levels without altering TERT expression (**Fig. 4C**), suggesting that these compounds may interfere with a distinct, noncanonical RdRP activity of TERT.

Collectively, these findings support the role of TERT as a putative RdRP contributing to the “msRNA → msdR → msdP” pathway. This expands the functional repertoire of TERT beyond telomere maintenance and implicates it in a novel regulatory axis relevant to cancer biology.

## Discussion

While prior research has centered mainly on the regulatory functions of msRNAs, particularly their roles in RNA silencing, our study reveals a fundamentally novel mechanism in human cells: specific pairing between the 3’-end of an msRNA and a target RNA can initiate the synthesis of new RNA molecules (msdRs), which in turn may be translated into functional polypeptides (msdPs). In this model, the msRNA and its paired RNA act as a “primer” and “template,” respectively—reminiscent of primer-dependent polymerase activity—thereby enabling RNA-dependent RNA synthesis in human cells. This discovery offers an expansion of the classical central dogma, deepening our understanding of RNA biology.

Even under a stringent 15-nt threshold, our analysis predicted over 1,000 msdPs, with an even greater number of msdRs. These newly predicted molecules contribute to the growing complexity of gene regulation and molecular biology in human cells. As a proof-of-concept, we validated one of these candidates, msdP0188, demonstrating its role in promoting tumor growth and metastasis in lung and breast cancers. Moreover, targeting msdP0188 through genetic or pharmacological means suppressed tumor progression, underscoring the therapeutic potential of the “msRNA → msdR → msdP” axis.

While this study establishes a conceptual and experimental foundation, the complete landscape and biological roles of msdRs and msdPs remain largely unexplored. Due to practical constraints, we were unable to validate all predicted molecules or fully characterize their biological functions. Following the same experimental protocols, we carried out a preliminary analysis of ten additional msdPs longer than 80 amino acids and/or their corresponding msdRs (**Table S8; Table S9**). RT-PCR followed by Sanger sequencing identified msdR fragments in eight of them (**Table S10; Data S11**). Of the four msdPs for which pAbs were produced, msdP0032 was successfully detected; LC-MS/MS analysis confirmed four of its peptide fragments with a total coverage of 30% (35 out of 118 amino acids) (**Fig. S17**). Furthermore, the 5’ RACE experiment verified that the 5’-end of msdRs0032 precisely matched the *hsa-miR-6892-3p* sequence, supporting its direct derivation from this priming msRNA (**Fig. S18; Data S11**). Along with the results for msdP0188 and msdRs0188, these findings suggest that many of these molecules deserve thorough follow-up investigations.

A major unresolved question concerns the molecular mechanism of msdR synthesis in human cells. The identification of msdRs supports the existence of an endogenous RNA-dependent RNA polymerization pathway, consistent with emerging evidence of such activity in human cells ^27,33^. Our knockdown and pharmacological inhibition experiments suggest that TERT may catalyze msdR synthesis, though the mechanistic details of TERT’s RdRP-like function remain unclear. Moreover, our bioinformatic analysis identified additional candidate proteins, including endogenous retroviral proteins, that possess either a conserved D*x*DGD motif or *palm*-like subdomains. These proteins may also contribute to msdR biogenesis, either independently or in conjunction with TERT.

Another possibility is that human cells acquire RdRP activity transiently during viral infection. Could viral RdRPs be repurposed by host cells to generate msdRs? Might host msRNAs pair with viral RNAs, or vice versa, to form new msdRs with functional or immunological implications? These questions raise exicting prospects for host-pathogen interactions and merit further investigations.

Several other scientific questions have arisen from our discovery, which warrant further investigation. Here, we provide some discussions based on related studies. First, it remains unknown whether msdRs possess canonical RNA modifications, such as a 5’-cap or 3’-poly (A) tail, and how msdRs can be stabilized if these features are absent, as these modifications are known to enhance transcript stability, nuclear export, and translation efficiency. Interestingly, RNAs located in the cytoplasm sometimes appear to be more stable than those in the nucleus ^34,35^. In tumor cells, some RNAs lacking poly(A) tails, such as c-Myc RNA, can be surprisingly stable ^36^, suggesting alternative stabilizing mechanisms. Given that msdRs likely localize to the cytoplasm and that some appear to be sufficiently stable for translation, it is plausible that msdRs utilize noncanonical mechanisms to maintain their integrity. However, these remain speculative and warrant rigorous experimental validation.

Second, in eukaryotic cells, double-stranded RNAs (dsRNAs) are often interpreted as pathogen-associated molecular patterns, triggering immune responses via sensors such as protein kinase R. Emerging evidence indicates that endogenous dsRNAs can also activate innate immune pathways in the absence of infection ^37,38^. Whether msdR/mRNA hybrids are recognized by such immune sensors—and whether this leads to inflammation or immune modulation—is an important and currently unanswered question.

Third, a critical question is how msdR/mRNA duplexes are unwound to allow ribosomal access and translation of msdRs. RNA helicases are likely involved in this process. Candidate helicases include MOV10, RNA helicase A, and DDX3X, each of which is known to unwind RNA duplexes with varying efficiency and directionality ^39–42^. Identifying the precise helicases involved will be key to understanding how msdRs are made translationally competent.

Our current computational approach assumes IRES-mediated translation initiation for msdRs. However, the necessity and sufficiency of IRES elements in msdP translation remain unproven. Moreover, the empirical nature of our IRES and start codon predictions may lead to false positives or negatives. Future iterations of the pipeline should incorporate broader models of translation initiation, including noncanonical mechanisms, to enhance sensitivity and specificity.

To conclude, our study reports a novel biological pathway in human cells wherein msRNAs can serve as primers for the synthesis of msdRs, which are then potentially translated into functional msdPs. This “msRNA → msdR → msdP” axis represents a previously unrecognized layer of gene expression regulation and an extension to the classical central dogma. We predict the existence of many previously unannotated RNAs and polypeptides that may play crucial roles in health and disease. The findings not only open a new dimension of RNA biology but also present a promising frontier for therapeutic intervention and biomarker discovery in cancer and potentially other diseases.

## Methods

### Prediction of msdRs and msdPs

We retrieved a total of 2,656 human miRNAs from miRbase (https://www.mirbase.org/; accessed on December 20, 2020), the largest repository for miRNA sequences and annotations. After discarding 24 redundant miRNAs (**Table S1**), the remaining 2,632 unique miRNAs were used as example msRNAs to derive msdRs and msdPs.

The 3’-end, such as a 15-nt fragment, of each msRNA (U → T) was queried against the NCBI RefSeq RNA database ^12^ using BLASTN ^13^ to identify complementary RNA templates for generating msdRs: *blastn -task blastn-short -db db_refseq_rna -query query_15nt.fa -out out_15nt.xml -outfmt 5 -evalue 1 - gapopen 5 -gapextend 2 -reward 1 -penalty -3 -ungapped -max_target_seqs 10000 -strand minus -taxids 9606*

Given that the queries are short, the “*-task blastn-short*” option was utilized, employing default parameters “*-gapopen 5 -gapextend 2 -reward 1 -penalty -3*” for “*blastn-short*” tasks. To ensure alignment without gaps, the “*-ungapped*” option was applied, while restricting the search to reverse matches was achieved with “*-strand minus*”. Further restriction to human RNAs was enacted via the “*-taxids 9606*” parameter.

We enforced strict reverse base pairing at the last bases of the 3’-end of msRNA with the RNA target through the following conditions:

1. The “-ungapped” option ensures no gaps in the pairing between msRNA and RNA.
2. We elaborately adjusted e-value thresholds, setting them at specific values such as 1, 5, 20, 100, 500, and 1,000 for queries of 15, 14, 13, 12, 11, and 10 nt, respectively. For instance, when utilizing the parameters “*-gapopen 5 -gapextend 2 -reward 1 -penalty -3 -ungapped*,” an exact, ungapped 15-nt match yields a calculated e-value of 0.94. Therefore, we employed a slightly higher e-value cutoff of “*-evalue 1*” for 15-nt queries. Hits with mismatches for the 15-nt query result in e-values larger than 1, hence such hits are excluded.
3. We mandated that the *Hsp_identity* (found in the *out_15nt.xml* file) be exactly equal to the query length (e.g., 15-nt) to ensure no mispairing.
4. We stipulated that “*-strand minus*” and “*Hsp_hit-frame = −1*” and “*Hsp_query-frame = 1*” (found in the *out_15nt.xml* file) to ensure that the msRNA is in the 5’→3’ direction while the paired fragment of RNA hit in the 3’→5’ direction.

With these BLASTN parameter settings, a sufficiently high cutoff of “*-max_target_seqs 10000*” was established to generate RNA hits; however, this threshold was not surpassed for queries ranging from 10 to 15 nt.

For clarity, the following discussion employs the 15-nt match as an example. Relevant information for a hit, including the GenInfo Identifier (GI) number, NCBI Reference Sequence number, start and end positions of the match, and gene symbol, was documented.

The sequence of a hit RNA was retrieved by utilizing its GI number:

*blastdbcmd -entry gi -db db_refseq_rna*

The msdR sequence was produced through reverse transcription of the RNA hit from the end-matching position to its 5’-end. The number of msdRs derived from each msRNA varies substantially. For instance, certain msRNAs (e.g., *hsa-miR-99b-5p*, *hsa-miR-9985*, and *hsa-miR-9903*, to name a few) do not produce any msdR, while *hsa-miR-619-5p* alone generates 3,421 msdRs. In total, 12,953 msdRs were obtained from all 2,632 human msRNAs. Of these, 11,121 unique msdRs were utilized to make a database:

*mkblastdb -in db_uniq_msdR.fa -out db_uniq_msdR -dbtype nucl*

The unique msdRs were translated into proteins if the following criteria were met: the msdR possesses a predicted IRES, and a start codon is present within 100 nt downstream of the IRES. The first criterion was verified by comparing msdRs with the experimental IRESs sourced from IRESbase ^14^, which encompasses 554 viral IRESs, 691 human IRESs, and 83 IRESs from other eukaryotes. Due to the considerable variation in sequence length, the experimental IRESs were searched against the msdR database using distinct BLASTN parameters:

If an IRES has ≤15 nt:

*blastn -db db_uniq_msdR -query ires_le15.fa -out ires_le15.xml -outfmt 5 -evalue 1e5 -task blastn-short - max_target_seqs 2500000 -reward 1 -penalty -3 -gapopen 5 -gapextend 2 -ungapped -strand plus*

Else if an IRES has <50 nt:

*blastn -db db_uniq_msdR -query ires_lt50.fa -out ires_lt50.xml -outfmt 5 -evalue 1e5 -task blastn-short - max_target_seqs 2500000 -reward 1 -penalty -2 -gapopen 0 -gapextend 2 -strand plus*

Otherwise:

*blastn -db db_uniq_msdR -query ires_ge50.fa -out ires_ge50.xml -outfmt 5 -evalue 1e5 -task blastn - max_target_seqs 2500000 -reward 2 -penalty -3 -gapopen 5 -gapextend 2 -strand plus*

The “*-strand plus*” option confines the search to forward matches. For short IRESs (≤15 nt), a high gap opening penalty “*-gapopen 5*” and the parameter *“-ungapped*” were applied to prohibit gaps; “*-penalty - 3*” was employed to discourage mismatches. For IRES spanning (15, 50) nt, penalty scores were adjusted to permit more mismatches and gap openings. For IRES >50 nt, default parameters for a standard “*blastn*” task were utilized. In all scenarios, large cutoffs for e-value (“*-evalue 1e5*”) and the number of targets (“*- max_target_seqs 2500000*”, exceeding the number of msdRs) were utilized.

An IRES site was considered to exist if *coverage* ≥ 1 and *aligned identity* ≥ 1 for IRES queries of ≤15 nt, or if *coverage* ≥ 0.7 and *aligned identity* ≥ 0.75 for queries >15 nt. Here, *coverage* and *aligned identity* were defined as the ratio of *aligned length* to *query length* and *identity* to *aligned length*, respectively. These metrics (*aligned length* and *identity*) were extracted from the BLASTN output (in the *.xml* file).

The msdRs were subsequently translated according to the second criterion. One msdR may encompass multiple IRESs, and there may be multiple start codons following an IRES. All potential scenarios were taken into account during the translation process from msdR to msdP(s). In total, 5,369 msdPs with more than 10 amino acids were generated, out of which 1,239 are unique. We conducted a BLASTP ^13^ search against the NCBI nr protein database to ascertain whether these msdPs have counterparts in the human protein database with high similarity using distinct parameters:

If a msdP has ≤15 amino acids:

*blastp -db nr -query msdP_le15.fa -out msdP_le15.xml -outfmt 5 -evalue 1 -task blastp-short -taxids 9606*

Else if a msdP has <30 amino acids:

*blastp -db nr -query msdP_lt30.fa -out msdP_lt30.xml -outfmt 5 -evalue 1e-4 -task blastp-short -taxids 9606*

Otherwise:

*blastp -db nr -query msdP_ge30.fa -out msdP_ge30.xml -outfmt 5 -evalue 1e-4 -task blastp -taxids 9606*

In this context, “*high similarity*” is characterized by *coverage* ≥ 1 and *aligned identity* ≥ 0.9 for msdPs with ≤15 amino acids, or *coverage* ≥ 0.9 and *aligned identity* ≥ 0.9 otherwise (*coverage* and *aligned identity* are parameters defined as previously described).

The same procedures were carried out for matching thresholds of 10–14 nt.

### Estimation of the potential false discovery rate (FDR) for msdR prediction

To evaluate the reliability of our bioinformatic pipeline, we estimated the potential false discovery rates (FDRs) for msdR prediction for different matching lengths. FDR was calculated as the ratio of false positives (FP) to predicted positives (PP), in which the number of msdRs predicted using database-documented msRNAs was regarded as PP whereas the number of msdRs predicted using computer-simulated msRNAs as FP. We carried out computational simulations to estimate FDR ranges.

As we predicted msdRs and msdPs using miRBase miRNAs, it was reasonable to take the number of msdRs predicted with unique human miRNAs (n = 2,632) in miRBase as PP. Inspired by previous studies utilizing shuffled miRNAs to estimate the FDRs for miRNA target prediction ^16,17^, we calculated FP as the mean number of predicted msdRs using n = 9 sets of msRNA artifacts (n = 2,632 of each set) by shuffling the 3’- end *l* (*l* = 10–15) nt of each parental miRBase miRNA (**Fig. S1**). The 3’-end nucleotide composition was held constant during shuffling, and therefore the msRNA artifacts generated in this way would exhibit identical binding affinities compared to the parental miRNAs when bound to RNAs with their 3’-end *l* nt.

We further estimated FDR ranges using randomly generated msRNA artifacts (**Fig. S1**), and we found that this method yielded lower FDR for each matching length.

### Nomenclature of msdR and msdP identifiers and names

We developed a nomenclature for assigning identifiers to msdRs and msdPs, reflecting the nature of their derivation from the elongation of msRNAs along template RNAs. The msdR identifier is structured as “msdR.{sRNA name}.{GenInfo Identifier number of the RNA (gene symbol of the RNA)}.{the end matching position on RNA}”, where “msdR” is a prefix. The msdP identifiers can be similarly established by extending the msdR identifier with {start codon position on msdR}. However, we observed that a msdP can be fully delineated by the template RNA alongside the position on RNA corresponding to the start-codon position on msdR. Therefore, a msdP identifier is formulated as “msdP.{GenInfo Identifier number of the RNA (gene symbol of the RNA)}.{the msdR start codon-paired position on RNA}”, where “msdP” is a prefix. This method largely avoids using multiple msdR-based msdP identifiers for the same msdP encoded by msdRs that originated from different msRNAs bound to the same template RNA.

Despite the meaningfulness, the above-defined identifiers may still not be unique for some msdRs/msdP due to RNA variants from splicing. To facilitate data presentation, we also used sequential names to describe certain msdRs and msdPs (**Table S4; Table S5; Data S4**).

### Estimation of the presence of msdPs via proteomics database search

The sequences of 1,193 new msdPs were used to create a FASTA database, which was then compared to mass spectrometry (MS) proteomics data obtained from four human tumor cell lines (refer to the “*Mass Spectrometry*” section) using Proteome Discoverer 2.1 (Thermo Scientific) with the Sequest-HT search engine. The S/N (signal-to-noise ratio) threshold was set to 1.5, the maximum missed cleavage sites to 2, the minimum peptide length to 6, the maximum peptide length to 144, the precursor mass tolerance to 10 ppm, and the fragment mass tolerance to 0.6 Da, respectively. Default parameters were used for other settings. The comparison suggested the potential presence of 941 msdPs, identified through a scoring metric that considers protein sequence coverage, the number of peptide-spectrum matches (PSMs), and the Sequest-HT score (**Data S8**).

We also used the standalone PepQuery2 to search for peptides within the 1,193 msdPs against the PepQueryDB ^19^ database. PepQuery serves as a targeted peptide search engine for the identification or validation of both known and novel peptides of interest in available MS-based proteomics datasets. PepQueryDB constitutes a comprehensive database of indexed MS/MS spectra, encompassing 48 datasets, 25,405 raw MS files, and a total of 1,012,696,284 MS/MS spectra. Following the PepQuery2 instructions, we executed the command “*java -jar pepquery-2.0.2.jar -b all -db gencode:human -hc -o $outdir/ -i $prot -fast -cpu $ncpu -n 1000 -t protein*”, where *$prot* denotes the query msdP protein sequence, *$outdir*specifies the output path, and *$ncpu* represents the number of CPUs employed for computation, respectively. The *“-t protein*” option specifies the input sequence as a digestible protein. The “*-n 1000*” parameter generates 1,000 random peptides to estimate the statistical significance of a PSM hit. The “*- fast*” option specifies the fast mode for searching. Confident PSMs were identified for 1,095 peptides originating from 555 msdPs (**Data S9**). Among them, peptides from 459 msdPs were also detected by the Sequest-HT search. Two peptide fragments that exhibit confident PSMs were discovered for msdP0188: LLETEVSLLSCGPALSCPCGVPAWGLHR and CLPRLLPSSWAPVGLLAFLWAR; notably, the latter encompasses the peptide LLPSSWAPVGLLAFLW determined by our MS analysis for verifying msdP0188. Similarly, two peptides featuring confident PSMs were identified for msdP0112: HPVFSLK and MDFSTKHPVFSLK.

### Prediction of msdR-deriving RdRP in human cells

We conducted a search of the human proteome (https://www.uniprot.org/proteomes/UP000005640, accessed on May 19, 2023), which encompasses 82,492 proteins associated with approximately 20,000 genes ^43^. Within this dataset, we considered proteins containing a linear D*x*DGD motif (*x* can be any amino acid) and consisting of >500 amino acids, as we reason that human RdRP-like proteins tend to have a relatively larger size. Subsequently, we manually analyzed these proteins, considering factors such as their function, subcellular location, structural information derived from experimental studies or predictions by AlphaFold2, as well as other features annotated in UniProt (**Data S12**).

On the other hand, we posited that it supplements the D*x*DGD motif-searching method by leveraging the *palm* subdomains of three viruses (SARS-CoV-2, human picobirnavirus strain Hy005102, and *Thosea asigna* virus) to potentially identify human RdRP-like proteins that lack the linear, signature D*x*DGD motif. We employed TM-align to explore structural analogs of the three viral *palm* subdomains across the entire Protein Data Bank (PDB) ^24^ database (204,826 structures; accessed on May 19, 2023) and the AlphaFold Protein Structure Database ^25,26^ for *Homo sapiens* (20,504 non-redundant models; accessed on May 19, 2023), utilizing a cutoff of TM-score > 0.5 (**Data S13; Data S14**).

### Cell line and transfection

The lung cancer cell line A549 and breast cancer cell line MDA-MB-231 were purchased from the Cell Resource Center of the Chinese Academy of Medical Sciences and authenticated through short tandem repeat analysis. All cell lines tested negative for mycoplasma contamination. A549 cells were cultured in minimum Eagle’s medium with 10% heat-inactivated fetal calf serum (A5669801, Gibco, USA) and supplemented with penicillin-streptomycin (C0222, Beyotime, China). MDA-MB-231 cells were cultured in Dulbecco’s modified Eagle’s medium with 10% heat-inactivated fetal calf serum (A5669801, Gibco, USA) and supplemented with penicillin-streptomycin (C0222, Beyotime, China). Cells were maintained at 37 °C with 5% CO_2_.

For the knockdown of msdP0188, shRNA lentivirus was synthesized by Obio Technology (Shanghai, China). RNA interference oligonucleotides were annealed and inserted into the pSLenti-U6-shRNA-CMV-EGFP-F2A-Puro-WPRE lentiviral vector. The interfering sequences are as follows: 5’-GGAGTCATCTCCACTGCATTT-3’ for shRNA-1, 5’-GCGTCTACTTCCTTCGTCTTG-3’ for shRNA-2, 5’-GCTAGGTGCGTCTGCTTCATT-3’ for shRNA-3, and 5’-CCTAAGGTTAAGTCGCCCTCG-3’ for control shRNA.

For the overexpression of msdP0188, the msdP0188-encoding sequence was cloned into PiggyBac Dual promoter to generate msdP0188-PiggyBac Dual promoter (GFP-Puro) plasmid by Genewiz Biotechnology (Beijing, China). Following the instructions provided by Neofect (Neofect Biotech Co., Ltd), the msdP0188-PiggyBac Dual promoter (GFP-Puro) plasmid and PiggyBac transposase (GFP-Puro) were transfected into A549 cells. After 48 hours, puromycin (1 μg/ml) was added to the culture for positive clone selection over the next 7 days. Confirmation of msdP0188-overexpressed A549 cell lines was conducted through Western blot analysis.

### Transwell assay

Transwell chambers (Corning, USA) were placed in a 24-well plate and 300 μl pre-warmed serum-free medium was added to the upper chamber. After incubating for 15 minutes at room temperature, the medium was discarded. The treated cells were then trypsinized into single cells and resuspended in a serum-free medium. After counting, the cells were seeded into the upper chamber at a density of 10^5^ cells/well. In the lower chamber, 500 μl of medium containing 10% fetal bovine serum medium was added. All cells were cultured in an incubator with CO_2_ at 37 °C for 24 hours. After the removal of cells from the upper chamber, the remaining cells were fixed with 4% formaldehyde for 30 minutes. Subsequently, the cells were washed with phosphate-buffered saline (PBS) and stained with 0.1% crystal violet. The invaded cells were then observed under a microscope at 200× magnification. All tests were done in triplicate.

### Wound healing assay

Cells were digested and seeded in 24-well plates at a density of 2×10^5^ cells/well. Upon cell density reaching 90%, straight scratches were made using a 100 μl pipette tip. The culture medium was then discarded, and cells were washed three times with PBS to eliminate debris. Subsequently, cells were cultured in serum-free medium in an incubator with 5% CO_2_ at 37 °C. After 48 hours, photographs were captured, and cell migration rates were calculated using the formula: (*initial wound distance* - *wound distance at 48 hours*) / *initial wound distance*. All tests were done in triplicate.

### Scanning electron microscope imaging

The cells were first fixed in pre-chilled 2.5% glutaraldehyde at 4 °C for 2 hours, followed by dehydration with a gradually increasing concentration of ethanol (30%, 50%, 70%, 80%, 90%, 95%, and 100% v/v) for 10 minutes at each concentration. Following this, cells underwent treatment with a gradually increasing concentration of tert-butanol (30%, 50%, 70%, 80%, 90%, 95%, and 100% v/v) for 15 minutes at each concentration. After removing the tert-butanol under a low-temperature vacuum, the cell surface was coated with gold. Finally, cell phenotypes were observed under a scanning electron microscope (JEOL, Japan).

### msdR RT-PCR

Total cellular RNAs were extracted using trizol (R0016, Beyotime, China) following the manufacturer’s instructions. The extracted total RNAs underwent treatment with DNase I (2270A, Takara, Japan) at 37 °C for 30 minutes to eliminate genomic DNA. Subsequently, 3 µg of total RNA was used for reverse transcription (RT) to obtain cDNA using the PrimeScript RT-PCR Kit (RR014A, Takara, Japan), employing msdR-specific RT primers rather than oligo (dT) or random primers. These RT primers, tailored to msdRs, were employed to minimize non-specific binding. Next, primers spanning at least two exons were designed for PCR amplification with the cDNA as a template. Amplification was carried out using the PrimeScript RT-PCR Reagent Kit (RR014A6110A, Takara, Japan) through 30 cycles at 94 °C for 30 seconds, 53 °C for 30 seconds, and 72 °C for 60 seconds. The PCR products were subsequently analyzed via 1% agarose gel electrophoresis and confirmed by DNA sequencing. All primer sequences are provided in **Table S6**.

### msRNA RT-qPCR

For msRNA RT-qPCR, total cellular RNAs were extracted using trizol (R0016, Beyotime, China) following the manufacturer’s instructions. 3 µg of total RNA was utilized for reverse transcription (RT) to obtain cDNA using the PrimeScript RT-PCR Kit (RR014A, Takara, Japan), employing msRNA-specific RT primers. U6 snRNA was used as an internal control. The resulting RT product was subsequently used as a template for quantitative real-time PCR using SYBR Green qPCR Mix (D7260, Beyotime, China) according to the manufacturer’s instructions. The ΔΔCt method was used to analyze the expression of msRNAs. All primers used in msRNA RT-qPCR are provided in **Table S11**.

### 5’ RACE of msdRs0188 and msdRs0032

Total RNA was extracted from A549 cells using the RNeasy Mini Kit (74106, QIAGEN, USA) following the manufacturer’s instructions. After incubation with DNase I to remove DNA contamination, 5 μg of total RNA was used to generate cDNA according to the protocol provided with the SuperScript™ III First-Strand Synthesis Kit (Thermo Fisher, USA). 5’-end phosphorylated sequence-specific RT primers for msdRs0188 (5’-pGGACGGATTACAGGCATGAGCCAAAAGGCGCAACTGAAGTA-3’) and msdRs0032 (5’-GAAATCTCCCACCCCTTGCAGCTGGAGTTGAAAGCCGACG-3’) were used for cDNA synthesis. The cDNA was designed to contain a terminal hybridization with four dangling nt at each end. To facilitate the terminal hybridization, cDNA was annealed from 90 °C to 22 °C at a rate of 0.1 °C/sec in 0.5× CircLigase buffer. Following this, the ends of annealed cDNA were ligated using single-strand DNA CircLigase (BIOSEARCH, USA) in 1× CircLigase buffer at 60 °C for 2 hours, resulting in fully closed single-stranded circular DNA (cssDNA). PCR was then performed to amplify the DNA fragment spanning the junction site of the cssDNA. Finally, Sanger sequencing was employed to identify the 5’-end of msdRs0188 and msdRs0032. PCR primers used are as follows: msdRs0188 forward primer, 5’-GAGGCAGAGGTTGTGGTGA-3’; msdRs0188 reverse primer, 5’-TCGTGGACTTTAGCCGAAT-3’; msdRs0032 forward primer, 5’-AGGCACCATCAGGAGCAG-3’; msdRs0032 reverse primer, 5’-AATCAGGACGTCAAAA GTTTG-3’.

### Western blot

Cellular proteins were extracted using RIPA lysis buffer (composed of 50 mM Tris, 150 mM NaCl, 1% NP-40, 0.5% sodium deoxycholate, and 0.1% SDS) containing protease inhibitors (P0013C, Beyotime, China). Following quantification with the BCA Protein Assay Kit (P0009, Beyotime, China), the proteins were separated via 10% SDS-PAGE and transferred onto a PVDF membrane (IPVH00010, Millipore, USA). The membrane was then immersed in QuickBlock blocking buffer (P0220, Beyotime, China) and subsequently incubated with the following antibodies for 2 hours at room temperature: anti-msdP0188 pAb (1:500), anti-TERT antibody (1:500, ABclonal, China), anti-tubulin antibody (1:1000, Affinity, USA), or beta-actin antibody (1:2000, Affinity, USA). Following this, the membranes were incubated with horseradish peroxidase-labeled secondary antibody (1:3000, Affinity, USA) at room temperature for 1 hour. After washing the membrane three times with PBS containing Tween20, protein bands were detected using the ECL Chemiluminescence Kit (P0018S, Beyotime, China). All tests were done in triplicate.

### Human telomerase reverse transcriptase (TERT) inhibition and knockdown

Two TERT inhibitors, BIBR-1532 (S1186, Sellect, USA) and zidovudine (HY-17413, MedChemExpress, USA), were used in the TERT inhibition assay. A549 cells were seeded in 24-well plates at a density of 2×10^5^ cells/well. After cell density reached 70%, they were treated individually with each inhibitor for specific durations and concentrations: BIBR1532 for 24 hours at 20, 50, 80, and 100 μM, respectively; and zidovudine for 60 hours at 20, 50, 80, and 100 μM, respectively. Untreated cells and dimethyl sulfoxide (DMSO) treated cells were used as controls.

Alternatively, TERT was knocked down using siRNAs, including two reported and three self-designed siRNAs. The sequences for indicated siRNAs were as follows: 5’-GAACUUCCCUGUAGAAGACGATT-3’ for siRNA-1 ^31^, 5’-GCAUUGGAAUCAGACAGCATT-3’ for siRNA-2 ^44^, 5’-GCGACGACGUGCUGGUUCACCTT-3’ for siRNA-432, 5’-CGGUGUACGCCGAGACCAAGCTT-3’ for siRNA-966, 5’-GGAAGAGUGUCUGGAGCAAGUTT-3’ for siRNA-1728, and 5’-UUCUCCGAACGUGUCACGUTT-3’ for control siRNA. A549 cells seeded in 24-well plates were transfected with siRNAs at a final concentration of 50 nM using Xfec RNA Transfection Reagent (631450, Takara, Japan). After 72 hours, cells were collected for protein extraction. Total proteins were extracted using the RIPA lysis buffer containing protease inhibitors (P0013C, Beyotime, China). Following quantification using the BCA Protein Assay Kit (P0009, Beyotime, China), proteins were separated by 10% SDS-PAGE and transferred to a PVDF membrane (IPVH00010, Millipore, USA). The membrane was then placed in 5% bovine serum albumin and incubated with the following antibodies: anti-msdP0188 pAb (1:500), anti-TERT antibody (1:500, ABclonal, China), or anti-tubulin antibody (1:1000, Affinity, USA), for 2 hours at room temperature, followed by incubation with horseradish peroxidase-labeled secondary antibody (1:3000, Affinity, USA) at room temperature for 1 hour. After washing the membrane three times with PBS-containing Tween20, protein detection was performed using the ECL Chemiluminescence Kit (P0018S, Beyotime, China). All tests were done in triplicate.

### Silver staining

After washing the cells twice with 1× PBS, proteins were extracted using 500 μl cell lysate (P0013C, Beyotime, China). A total of 50 µl protein A/G magnetic beads (HY-K0202, MCE, USA) were then incubated with the anti-msdP0188 pAb or rabbit IgG (negative control) at room temperature for 30 minutes. Subsequently, the harvested beads were incubated with the extracted proteins at 4 °C for 2 hours. After elution using 1× SDS-PAGE loading buffer, the protein complex was separated by 12% SDS-PAGE electrophoresis. Silver staining was performed following the provided instructions (P0017S, Beyotime, China). The gel was fixed with 100 ml of fixative solution at room temperature for 20 minutes, followed by washing with 30% ethanol and pure water for 10 minutes each. The silver solution was then applied for 10 minutes at room temperature, followed by washing with pure water. The reaction was terminated by treatment with the color-developing solution for 10 minutes. After photography, the ∼18 kDa protein band of interest was excised for subsequent analysis.

### Mass spectrometry (MS)

#### Generation of MS spectra data from human tumor cell lines

After harvesting, the four tumor cell lines (HeLa, PANC-1, U118, and HCT116) underwent a triple wash with 1× PBS followed by the addition of RIPA buffer (R0010, Solarbio, China) containing a mixture of protease inhibitors (P6730, Solarbio, China) to induce cell lysis. After centrifugation, the resulting supernatant was subjected to protein concentration measurement using a BCA kit (PC0020, Solarbio, China). Subsequently, 5× loading buffer (P1040, Solarbio, China) was added to the cell lysate, which was then boiled for 10 minutes. Following this, 100 μg of protein was subjected to separation via 10% SDS-PAGE electrophoresis. The gel was then stained with Coomassie brilliant blue R250 solution for 20 minutes, and the color was removed using a decolorization solution comprising 50 mM NH_4_HCO_3_ and acetonitrile (1:1, v/v) until the background was clear. The gel was cut into approximately 1-mm^3^ pieces, and decolorization continued until the blue coloration was completely faded, followed by dehydration with acetonitrile. Reduction alkylation of proteins was achieved by sequential addition of 10 mM DTT (treated at 56 °C for 1 hour) and 55 mM IAA (treated at room temperature in the dark for 45 minutes) to the gel. The gel was then rinsed twice with 25 mM NH_4_HCO_3_ and a mixture of acetonitrile and 50 mM NH_4_HCO_3_ (1:1, v/v), respectively, before being dehydrated again with acetonitrile. Trypsin was then added, and the reaction proceeded for 16 hours at 37 °C. The resulting digested peptides were extracted with a solution of acetonitrile, formic acid, and water (7:1:2, v/v/v), and subsequently dried using a vacuum concentrator. The peptides were redissolved in an aqueous solution of 0.1% formic acid for liquid chromatography with tandem mass spectrometry (LC-MS/MS) detection. The LC-MS/MS system comprises an EASY-nLC system and an Orbitrap Fusion Tribrid mass spectrometer. Separation was conducted using an Acclaim PepMap 100 column (75 µm × 15 cm, nanoViper, C18, 3 µm, 100 Å; Thermo Scientific), with a mobile phase consisting of acetonitrile (B) and 0.1% formic acid aqueous solution (A) at a flow rate of 300 nl/min. The composition of the mobile phase starts at 97% A, linearly decreasing with the following schedule: 97%–92% A (0–5 minutes), 92%–82% A (5–75 minutes), 82%–72% A (75–103 minutes), 72%–10% A (103–115 minutes), and finally keeping at 10% A for 5 minutes. MS data acquisition utilized a data-dependent acquisition (DDA) mode, with scanning and fragmentation modes of MS being orbitrap (OT)–higher energy collisional dissociation (HCD)–iontrap (IT).

#### Identification of msdP0188 and msdP0032 using MS

To identify msdP0188, we utilized an Acuity UPLC system (Thermo Scientific) coupled with a Q Exactive orbitrap mass spectrometer (Thermo Scientific) for protein identification through peptide mapping. Following silver staining of the SDS-PAGE gel, the ∼18 kDa band of interest was excised, de-stained, reduced, and alkylated. Subsequently, in situ trypsin digestion was performed for 16 hours at 37 °C. The resulting digested peptides were then extracted using a solution of acetonitrile, formic acid, and water (7:1:2, v/v/v), and dried directly via vacuum concentration. Peptide separation was achieved using a UPLC system (Thermo Scientific) equipped with a reversed-phase CSH C18 column (150 mm, 1.7 μm, Waters), with detection performed by a Q Exactive orbitrap mass spectrometer (Thermo Scientific) operating in high-sensitivity mode. The flow rate was set at 200 μl/min, with a column temperature maintained at 40 °C. Solvent A consisted of 0.1% formic acid (FA) in water, while solvent B consisted of 0.1% FA in acetonitrile. During the 120-minute gradient, the percentage of solvent B level was increased progressively: from 2% to 29% over 60 minutes, from 29% to 55 % in 5 minutes, from 55% to 75% in 2 minutes, and held at 75% for 5 minutes. Subsequently, it was returned to 2% over 2 minutes and maintained at 2% for the remaining 6 minutes. The digestion products by Glu-C (staphylococcal V8 protease) and chymotrypsin were similarly analyzed.

For msdP0032 identification, the ∼12 kDa band observed in the silver-stained SDS-PAGE gel was excised and processed for in-gel digestion and mass spectrometry analysis by MS Bioworks (Ann Arbor, MI, USA). Briefly, gel pieces were washed with 25 mM ammonium bicarbonate followed by acetonitrile, reduced with 10 mM dithiothreitol at 60 °C, and alkylated with 50 mM iodoacetamide at room temperature. Proteins were digested in situ with trypsin (Promega) at 37 °C for 4 hours, and the reaction was quenched with formic acid. Peptides were separated using a Waters M-Class nano-LC system coupled to a Thermo Scientific Exploris 480 Orbitrap mass spectrometer. Separation was performed on a 75 μm analytical column packed with XSelect CSH C18 resin (2.4 μm particle size, Waters) at a flow rate of 350 nl/min. The column temperature was maintained at 55 °C using a Sonation column heater. The mass spectrometer was operated in data-dependent acquisition mode, with MS and MS/MS resolutions set at 60,000 and 15,000 FWHM, respectively, using a 3-second duty cycle. Chymotrypsin digestion products were analyzed using the same workflow.

### Synthesis of Circular single-stranded DNA (CSSD)

CSSD was designed to contain multiple complementary antisense sequences for msRNA absorption. Here CSSD4 contains reverse complementary sequences of four msRNAs, while the control CSSD comprises four repetitive scrambled sequences (see below, underlined). All nucleic acids have phosphorylated 5’-ends.

For control CSSD, the single-stranded DNA 5’-AATTCAAAGAATTAACCTTAATTGAAGGGGAGGGTT<u>CAGTACTTTTGTGTAGTACAA</u>ATAT<u>CAGTACTTTTGTGTAGTACAA</u>AAGGGAGGGCTTCAATTAAGGTTAATTCTTTG-3’ was diluted in annealing buffer (Beyotime, China), heated at 95 °C for 2 minutes, and then cooled to 4 °C at a rate of 0.1 °C per 8 seconds. Subsequently, the annealed DNAs were treated with T4 DNA ligase (Takara, Beijing, China) for 2 hours at room temperature to form a complete circular DNA.

For CSSD4, two single-stranded DNAs were synthesized: 5’-AATTCAAAGAATTAACCTTAATTGAAGGGGAGGGTT<u>CACCACTGCACTCGCTCCAGCC</u>ATAT<u>GGCTCATGCCTGTAATCCCAGC</u>AAGGGAGGGCTTCAATTAAGGTTAATTCTTTG -3’ containing binding sequences that are reverse complementary to *miR-619-5p* and *miR-3135b* (underlined) and 5’-AATTCAAAGAATTAACCTTAATTGAAGGGGAGGGTT<u>CTGGGGCAGGAGGGAGGTCAGCATG</u>ATAT<u>ACCTGTAGTCCCAGCTATTCAG</u>AAGGGAGGGCTTCAATTAAGGTTAATTCTTT-3’ containing binding sequences that are reverse complementary to *miR-4728-3p* and *miR-5585-3p* (underlined). After separate annealing with annealing buffer (Takara, Beijing, China), the two DNA strands were treated with T4 DNA ligase (Takara, Beijing, China) to form a completely closed circular DNA (**Fig. S11**). All single-stranded DNAs were synthesized by Genewiz Biotechnology (Beijing, China).

### Immunohistochemistry (IHC) assay

Various tissue samples, including 15 lung cancer tissues, nine normal lung tissues, 10 estrogen receptor-positive (ER+) breast cancer tissues, 12 triple-negative breast cancer (TNBC) tissues, and eight normal breast tissues, were analyzed using IHC assays. Tumor tissues were fixed with 10% formalin and embedded in paraffin. For the IHC analysis, the embedded tissues were sectioned into four μm-thick slices. Antigen retrieval was performed using a sodium citrate solution (Beyotime, China), followed by blocking with 3% hydrogen peroxide. Subsequently, the sections were incubated overnight at 4 °C in the dark with anti-msdP pAb (1:200). Negative controls were prepared by omitting the anti-msdP pAb during the incubation step. After washing with PBS, the samples were incubated with a horseradish peroxidase-conjugated secondary antibody (1:2000, Abcam, UK). Subsequently, staining development was carried out using the DAB kit (Beyotime, China), followed by counterstaining with hematoxylin. As msdPs are novel proteins and their tissue-specific expression is not yet fully understood, establishing a positive control for IHC analysis is currently not feasible.

### Polyclonal Antibody (pAb)

For selected msdPs, specific immunogenic peptides were designed and synthesized (Apeptide, Shanghai, China) and confirmed through mass spectrometric analysis. These immunogenic peptides have no homologous sequences with the annotated proteins after blast comparison. An additional Cys residue was appended at the C-terminal for conjugation with the keyhole limpet hemocyanin protein. The immunogenic peptide sequences are provided in **Table S4**.

The immunogens were then used to immunize rabbits. In the initial immunization, 350 µg of antigen was mixed with 1,000 μl of Freund’s complete adjuvant (Sigma, F5881) and subcutaneously injected at multiple sites. Subsequently, for three follow-up immunizations, 200 µg of antigen mixed with 1,000 μl of Freund’s incomplete adjuvant (Sigma, F5506) was administered subcutaneously using the same procedure. The interval between immunizations was 14 days. On day 7 following the final immunization, the rabbits were euthanized, and blood was collected and centrifuged to obtain antiserum for subsequent antigen affinity purification. This method is widely used to get the purest pAbs with the least amount of cross-reactivity. The antiserum from immunized rabbits was incubated in a column filled with epoxy-activated agarose (Wechsler Chromatography, Beijing, China) conjugated with peptide immunogens, and then eluted to obtain relatively purified antigen-specific pAbs.

We demonstrated the pAbs’ sensitivity and specificity through a series of experiments, using anti-msdP0118 pAb as an example. We constructed a vector, pET28a-msdP0188, to overexpress msdP0188 in *E. coli* and obtain recombinant msdP0188. The purified msdP0188 protein was used as a positive control in Western blot analysis to demonstrate the sensitivity of the anti-msdP0188 pAb, as we expected the anti-msdP0188 pAb to detect msdP0188 effectively. SDS-PAGE was run to separate recombinant msdP0188 and A549 cell lysate simultaneously, and in all the groups, ∼18 kDa bands that match the calculated molecular weight were detected (**Fig. S4**). Following this, we conducted IHC staining by treating human lung cancer and triple-negative breast cancer tissues with anti-msdP0188 pAb, using horseradish peroxidase-labeled goat anti-rabbit secondary IgG (isotype IgG)-treated or untreated (PBS) human cancer tissues as negative controls (**Fig. S5**). Robust staining signals were observed in the anti-msdP0188 pAb group, whereas the two control groups showed no or much weaker discernible signals (**Fig. S5**). Besides, in the silver staining experiment we also detected a ∼18 kDa protein band in the A549 cell lysate using protein A/G magnetic beads that are capable of binding anti-msdP0188 pAb. We confirmed the identity of proteins within this band as msdP0188 through MS analysis (**Fig. 1**). Following the same bioinformatic pipeline, we did not anticipate the presence of msdP0188 in mice. As expected, in contrast to the robust signals observed for the anti-msdP0188 pAb in human cancer tissues (**Fig. S5**), IHC assays in mouse liver and spleen with anti-msdP0188 pAb or PBS showed no detectable staining signals (**Fig. S6**). Together, these data demonstrate the high sensitivity and specificity of the anti-msdP0188 pAb to target msdP0188.

### Clinical specimens

A pathologist reviewed tumor samples embedded in paraffin from the First Hospital Of Qinhuangdao. The study was conducted in accordance with the guidelines outlined in the Declaration of Helsinki and its subsequent amendments, or with comparable ethical standards.

### Xenograft tumor in nude mice

In the tumor growth experiments, 24 five-week-old BALB/C nude mice were randomly divided into four groups. Two groups were subcutaneously inoculated with A549 cells at a concentration of 5×10^6^ cells/mouse and subsequently treated with CSSD4 or control CSSD via intratumoral injection at a dosage of 20 μg/mouse, administered once every seven days for a total of three treatments. The same procedures were repeated with the other two groups using MDA-MB-231 cells. Tumor sizes were measured every three days, and tumor volume was calculated using the formula: *tumor volume = (major axis of tumor)×(minor axis of tumor)*^2^*/2*. At the end of the experiment, all mice were euthanized on day 34 via CO_2_ asphyxiation. Tumor tissues were fixed and embedded in paraffin for subsequent analysis. The expression of msdP0188 in solid tumors was detected using immunohistochemical staining and the bright-field imaging was captured using an Olympus microscope.

In the tumor metastasis experiments, 12 five-week-old BALB/C nude mice were randomly divided into four groups. Two groups of mice were inoculated with A549 cells via the tail vein at 5×10^6^ cells/mouse. Ten days later, CSSD4 or control CSSD was injected into the tail veins for treatment at a dosage of 30 μg/mouse, administered once every seven days for a total of three treatments.

All animal experiments were conducted in compliance with the regulations outlined by the Institutional Animal Care and Use Committee (IACUC) of the Tianjin International Joint Academy of Biomedicine.

### Statistical analysis

Statistical analyses were conducted using GraphPad Prism 9.0 (La Jolla, CA, USA), and results are presented as the mean ± s.e.m. Comparisons between two groups were performed using an unpaired, two-tailed Student’s *t*-test, while multiple-group comparisons were analyzed using one-way ANOVA followed by Tukey’s post-hoc test. \**P* < 0.05; \*\**P* < 0.01; ns, not significant.

## Data and code availability

The miRBase miRNA sequences were retrieved from https://www.mirbase.org/ftp.shtml (accessed on December 20, 2020). Experimentally determined IRES sequences were obtained from http://reprod.njmu.edu.cn/cgi-bin/iresbase/index.php (accessed on January 2, 2021). The NCBI RefSeq RNA database was acquired from https://ftp.ncbi.nlm.nih.gov/blast/db (accessed on December 20, 2020). The NCBI nr protein database was obtained from the same source (accessed on January 2, 2021). The BLAST suite (version 2.11.0) was downloaded from https://ftp.ncbi.nlm.nih.gov/blast/executables/blast+. The human proteome was retrieved from https://www.uniprot.org/proteomes/UP000005640 (accessed on May 19, 2023). The Protein Data Bank (PDB) was accessed from https://files.rcsb.org/pub/pdb/data/structures/divided/mmCIF (accessed on May 19, 2023). The AlphaFold Protein Structure Database (AFDB) for Homo sapiens was accessed from https://alphafold.ebi.ac.uk/download (accessed on May 19, 2023). The PepQuery2 program (version 2.0.2) was downloaded from http://pepquery.org/download.html (accessed on December 7, 2023). The PepQueryDB database is available at http://pepquery.org/index.html (accessed by PepQuery2 with Internet connection). In case these databases or software have been updated, the specific versions employed in this study can be provided upon request. All scripts for predicting and analyzing msdRs and msdPs, along with supplementary data and other related source data, are made publicly available at https://doi.org/10.5281/zenodo.7826531.

## Acknowledgments

We thank the Advanced Research Computing (ARC) at the University of Michigan for providing the computational resources and services (to X.H.) that supported this research.

## Author contributions

X.H., H.W., J.X., Y.E.C., and S.C. conceived the project. X.H., Y.E.C., and S.C. supervised and administrated the project. X.H. conducted all computational research. H.W., W.L., M.Z., X.K., R.Z., K.L., X.L., Y.L., Z.Y., M.W., and S.C. conducted transwell, wound healing, and SEM experiments. X.X. and H.Z. performed the MS experiment. Z.L., W.Z., B.C., Z.Y., and S.C. conducted the animal experiment. W.Z., X.L., and Z.L. conducted the IHC experiment. H.W., W.Z., W.L., R.Z., Z.Y., X.W., Y.X., and S.C. carried out PCR and Western blot experiments. H.W., G.Z., Z.W., and S.C. conducted the silver staining experiment. J.Zhong., Z.Y., and S.C. performed the 5’ RACE experiments. H.W., W.L., Z.Y., and Z.L. designed msdP-specific antigens and produced antibodies. X.H., H.W., W.Z., Z.L., X.W., Y.H., Y.P., and S.C. performed formal analysis. X.H., Z.Y., M.Z., M.W., G.Z., Y.H., J.X., Y.E.C., and S.C. provided resources. X.H. and S.C. wrote the initial manuscript. X.H., J.Zhang, J.X., Y.E.C., S.C., Z.Y., and G.S.O. edited the manuscript. All authors participated in discussions and approved the final manuscript.

## Supplementary Materials

**Fig. S1.**
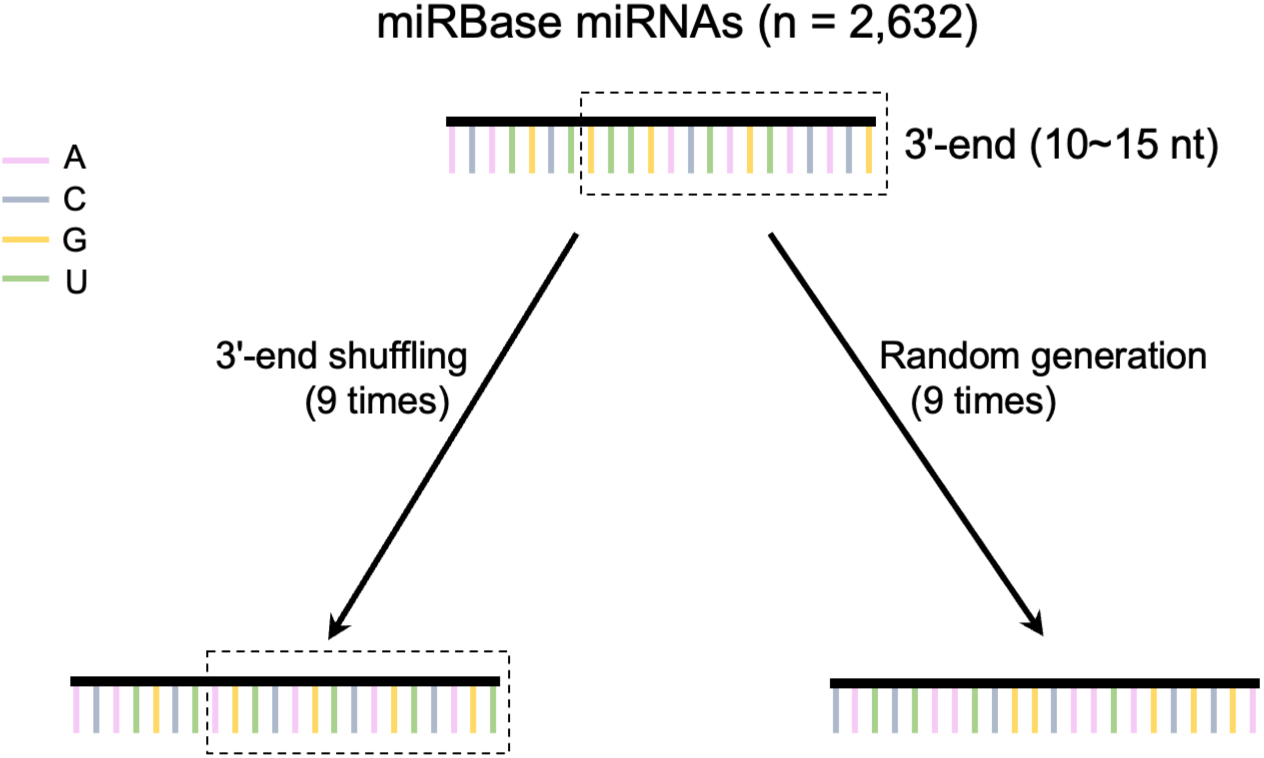
Schematic representation of methods used to generate control msRNAs for estimating the false discovery rate in msdR prediction. Techniques including 3’-end shuffling and random sequence generation based on unique human miRNAs from miRBase (n = 2,632).

**Fig. S2.**
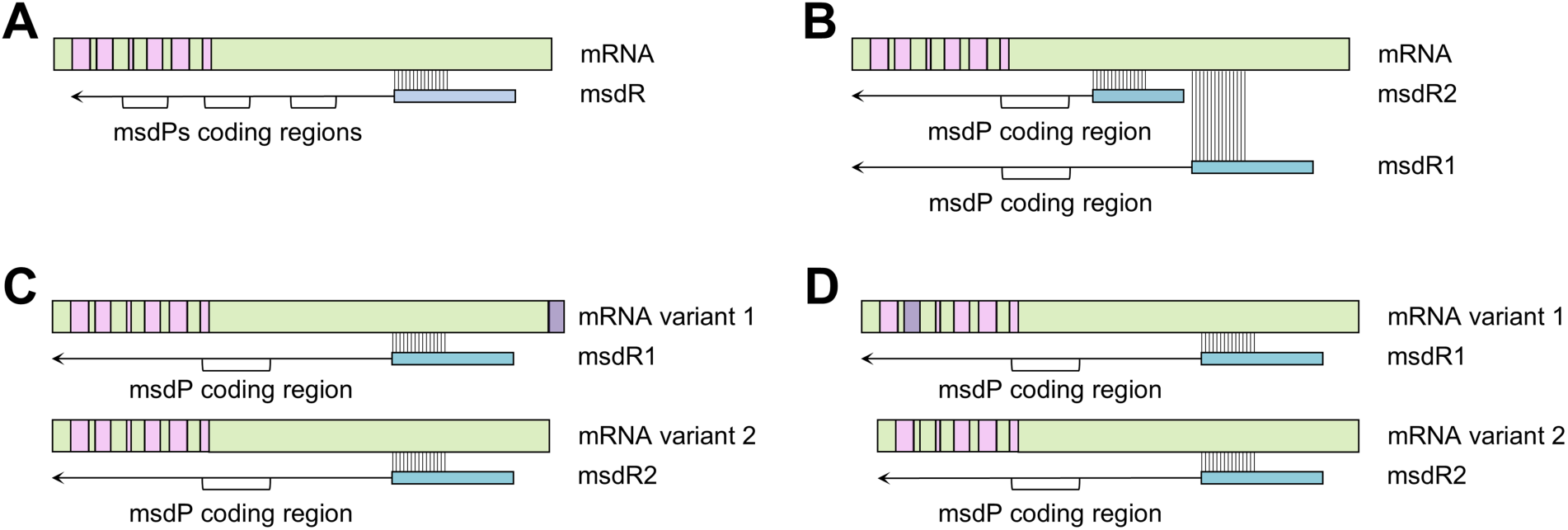
Possible scenarios for msdR and msdP derivation. (**A**) A single msdR can encode multiple msdPs. **(B)** Different msRNAs binding to the same RNA yield distinct msdRs, which may encode identical msdPs. **(C)** RNA splicing outside the msRNA elongation region results in identical msdRs. (**D**) RNA splicing within the msRNA elongation region produces different msdRs, though they may encode identical msdPs. In (C) and (D), the excised exon in mRNA variant 1 is depicted in purple.

**Fig. S3.**
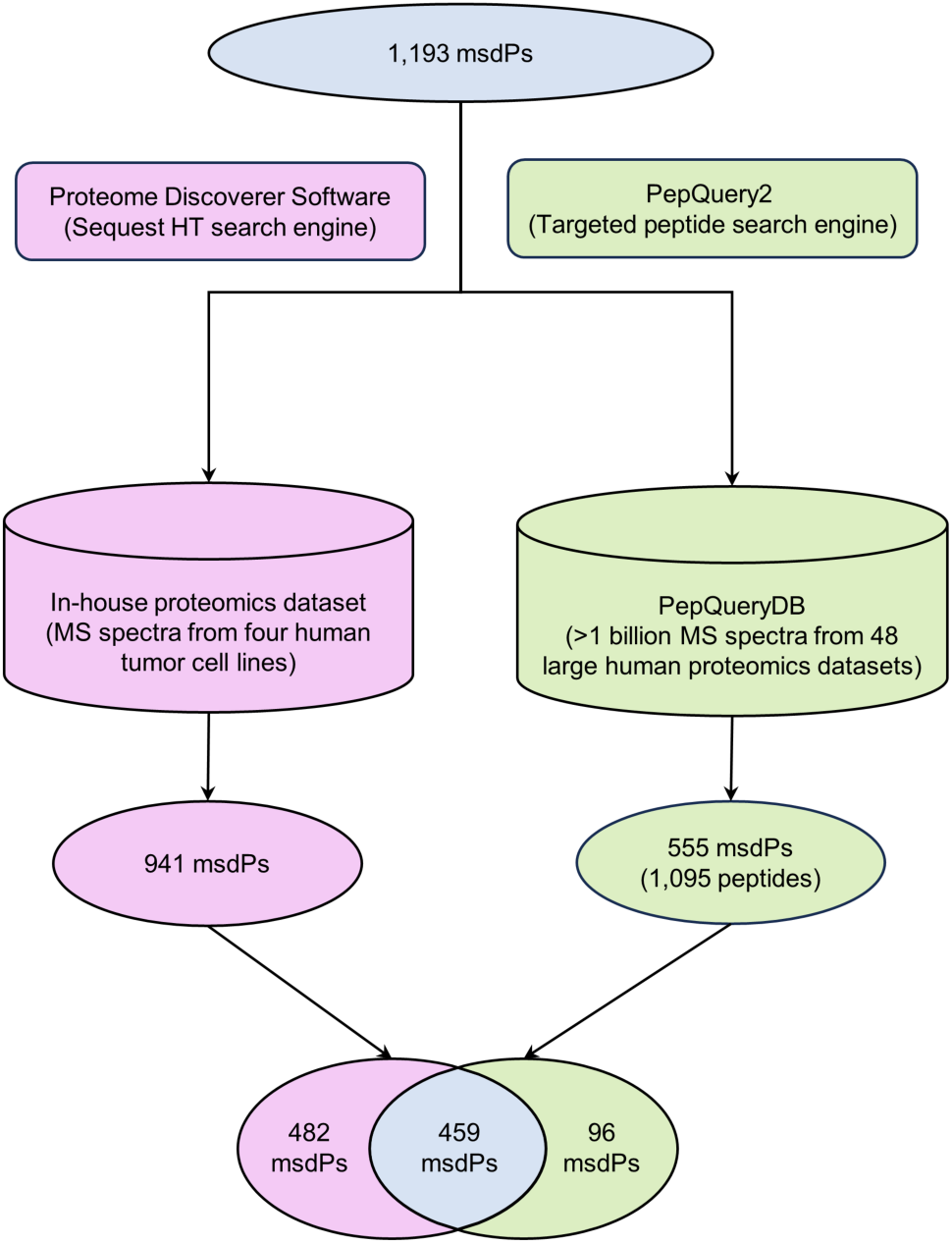
Workflow for searching msdP peptide fragments in in-house and public proteomics databases. MS, mass spectrometry.

**Fig. S4.**
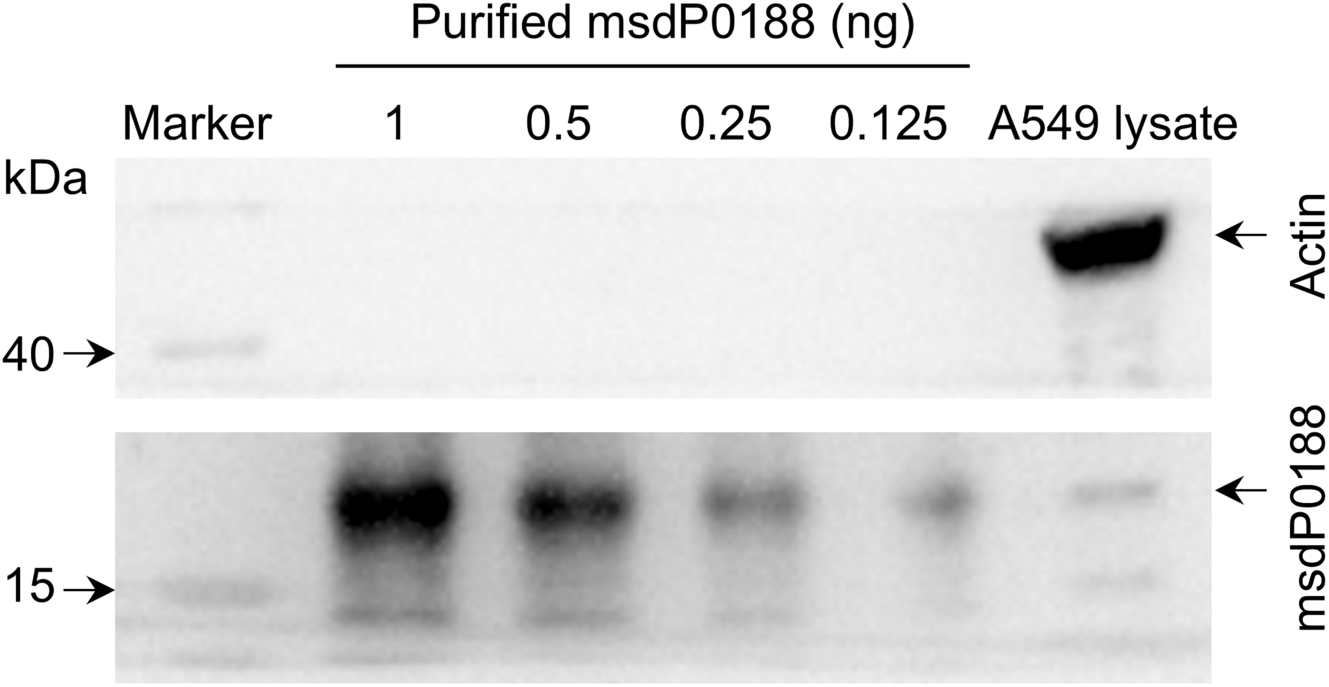
Western blot analysis of purified msdP0188 protein and A549 cell lysate using the anti-msdP0188 polyclonal antibody. Serial dilutions of purified msdP0188 were loaded for analysis.

**Fig. S5.**
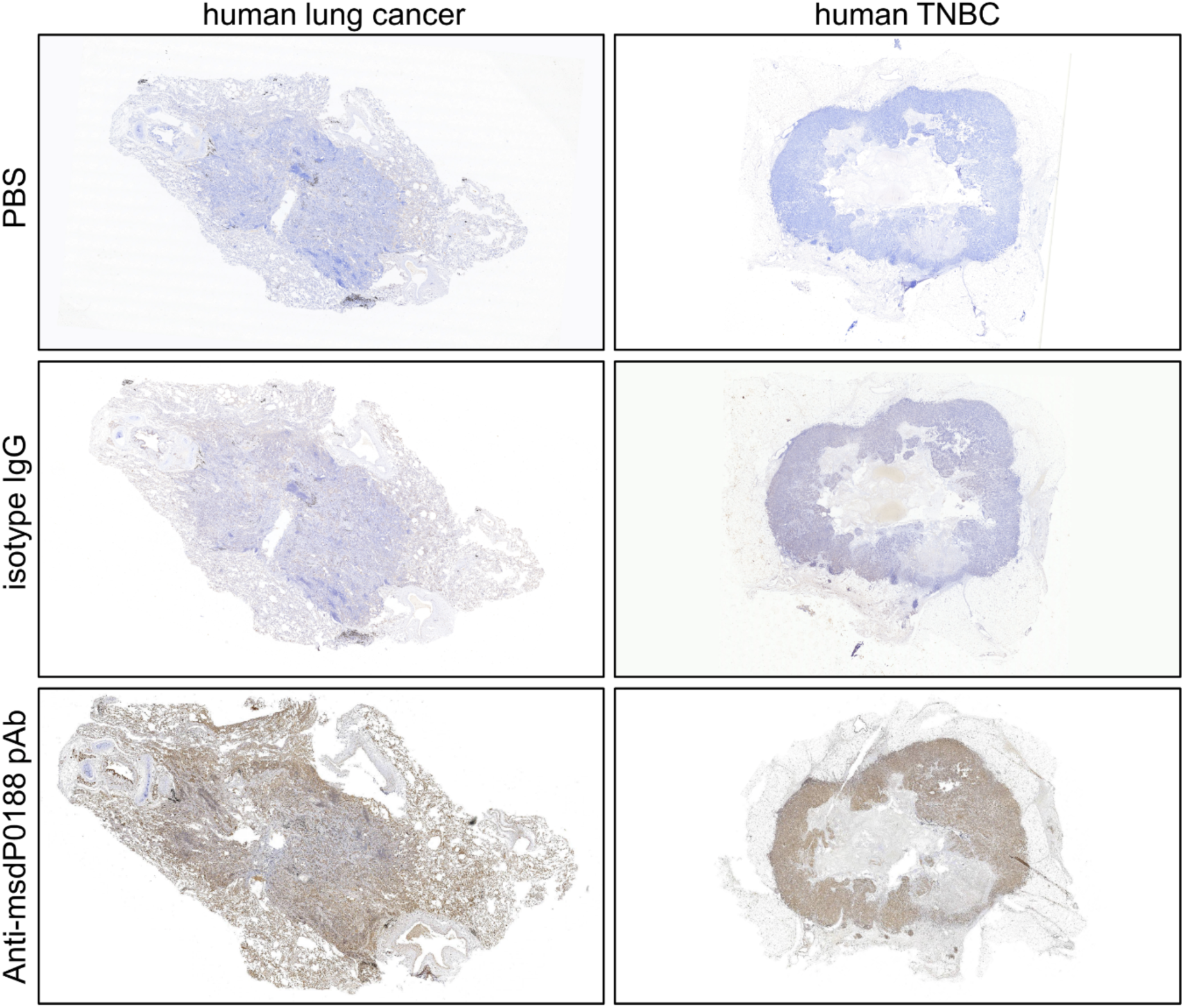
Immunohistochemical staining of human cancer tissues. pAb, polyclonal antibody. PBS, phosphate-buffered saline. TNBC, triple-negative breast cancer.

**Fig. S6.**
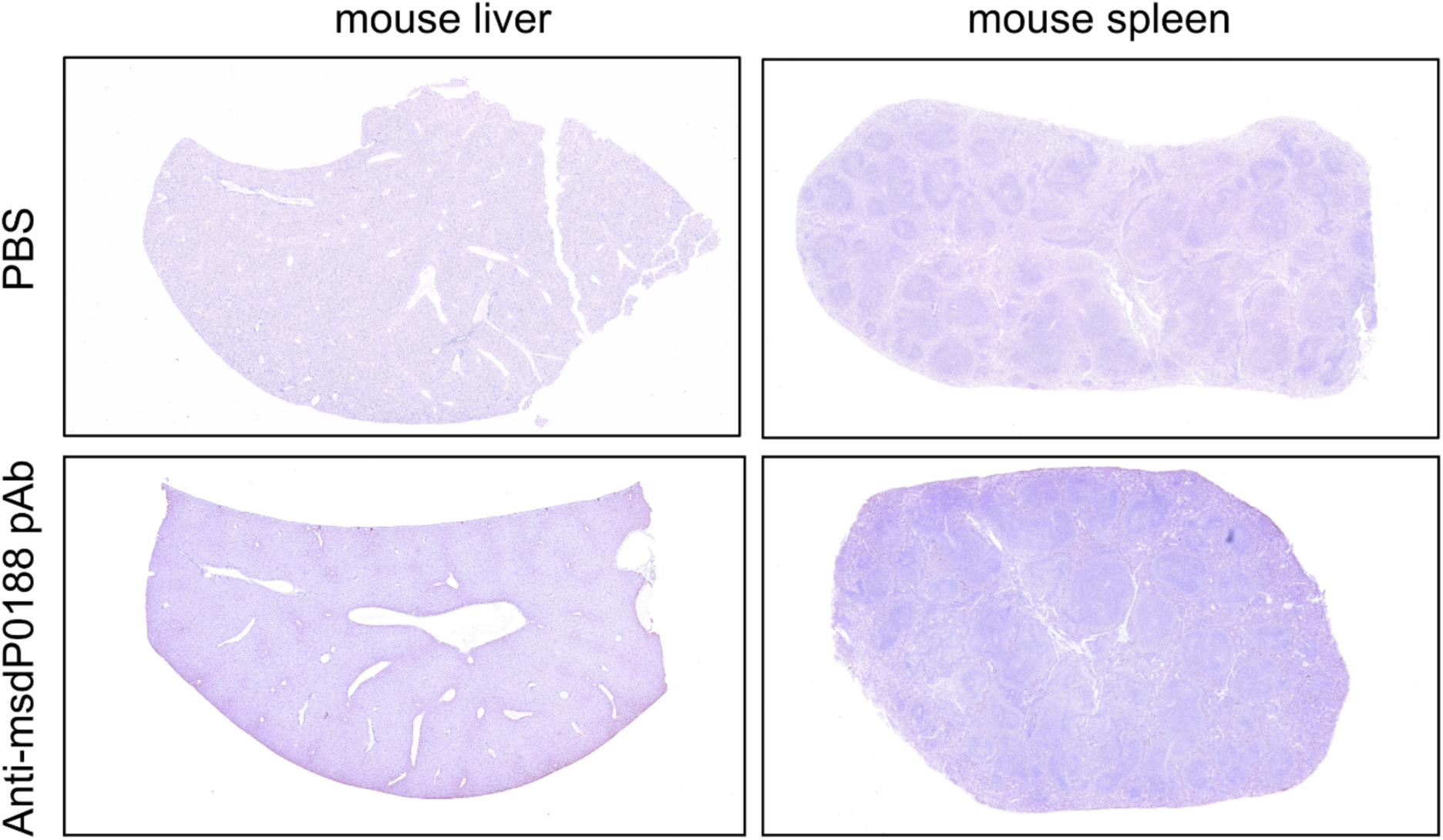
Immunohistochemical staining of mouse tissues. pAb, polyclonal antibody. PBS, phosphate-buffered saline.

**Fig. S7.**
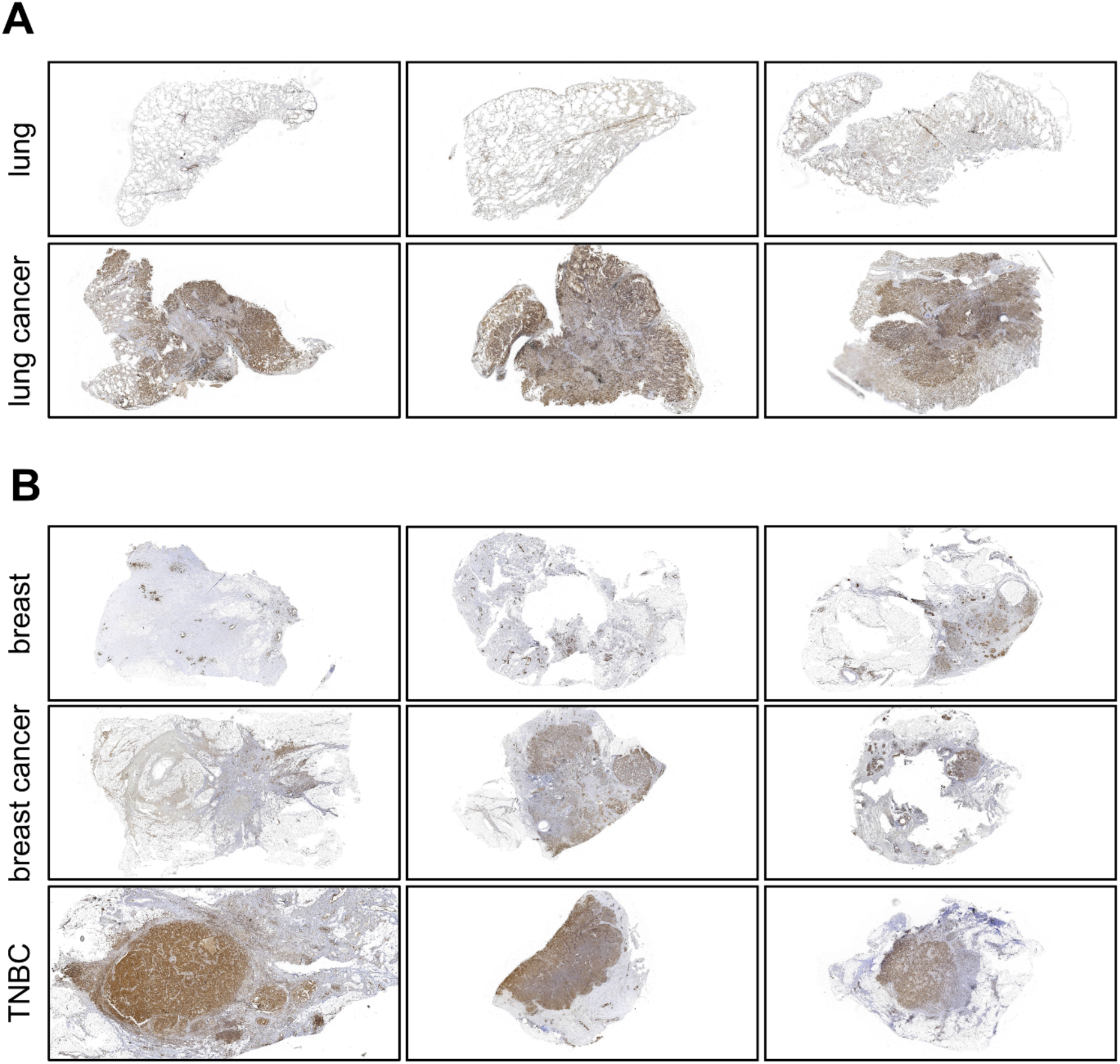
Immunohistochemical staining of tissues from various patients. (**A**) Full-section images stained with anti-msdP0188 polyclonal antibody (pAb) showing normal lung tissues (n = 3) and lung cancer tissues (n = 3). (**B**) Full-section images stained with anti-msdP0188 pAb stained full-section images showing normal breast tissues (n = 3), breast cancer tissues (n = 3), and triple-negative breast cancer (TNBC) tissues (n = 3). Each image corresponds to a different patient.

**Fig. S8.**
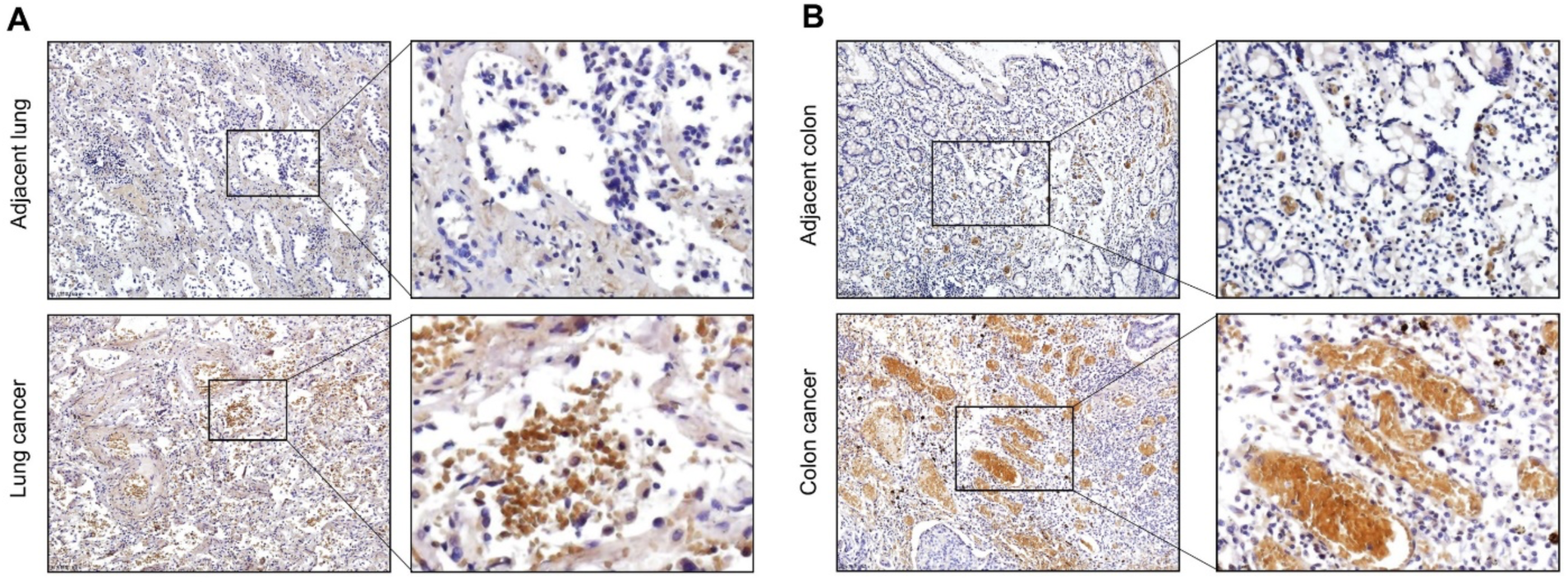
Expression of msdP0112 in lung and colon cancer tissues. (**A**) Immunohistochemical staining of lung cancer and adjacent lung tissues using the anti-msdP0112 polyclonal antibody (left). Sections enclosed by black rectangles are magnified to 100× (right) for each core section. (**B**) Immunohistochemical staining of colon cancer and adjacent colon tissues using the anti-msdP0112 polyclonal antibody (left). Sections enclosed by black rectangles are magnified to 100× (right) for each core section.

**Fig. S9.**
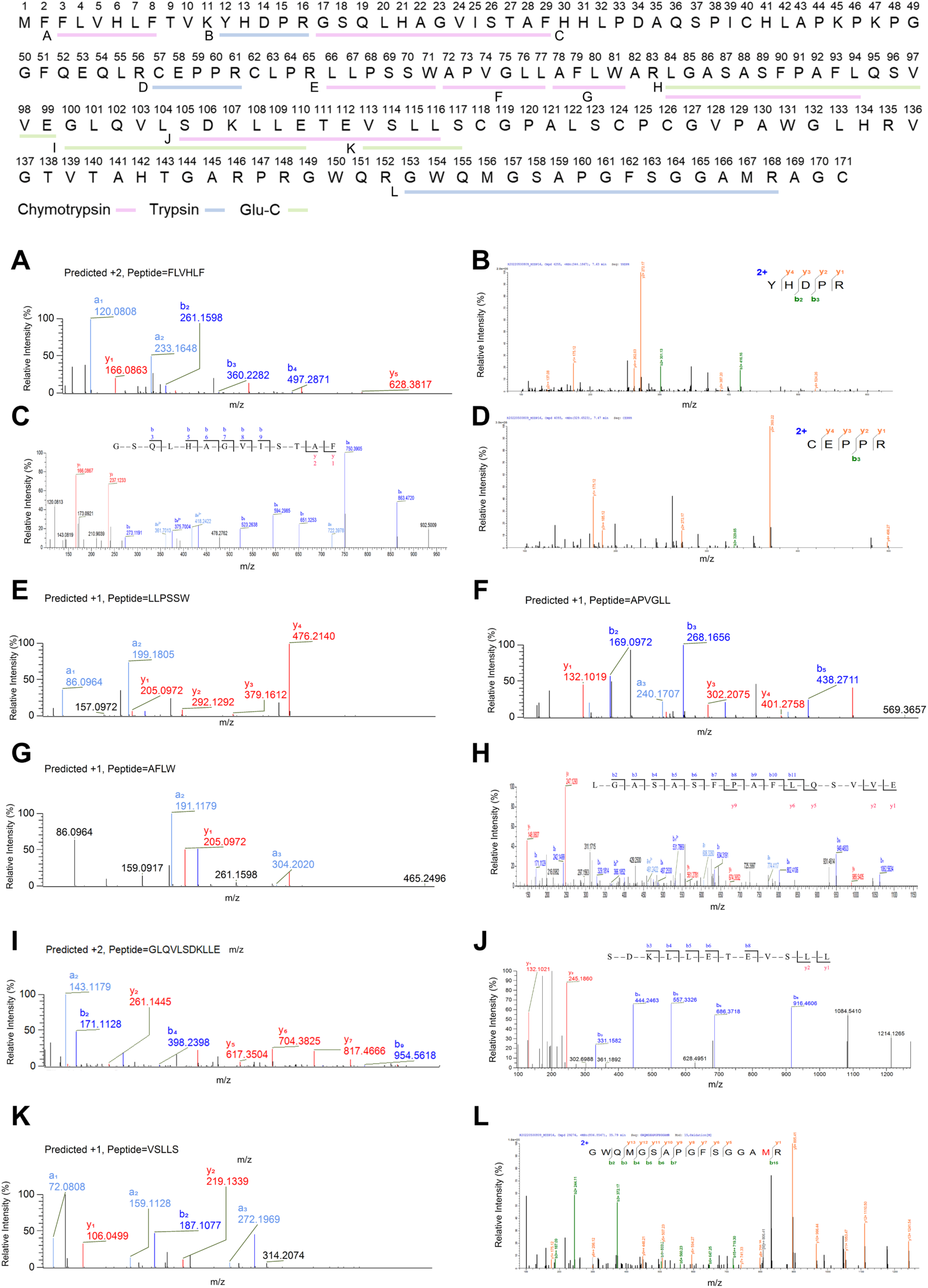
Mass spectrometry assay authenticates msdP0188 peptide fragments after protease digestion. The protein bound to the anti-msdP0188 polyclonal antibody during immunoprecipitation was separated via SDS-PAGE and silver staining to identify a potential target band. This band was isolated and subjected to digestion individually with three proteases. The digested samples were then analyzed using mass spectrometry. Thirteen peptide fragments within msdP0188 were validated, including three from trypsin digestion (blue), three from Glu-C digestion (green), and seven from chymotrypsin digestion (pink).

**Fig. S10.**
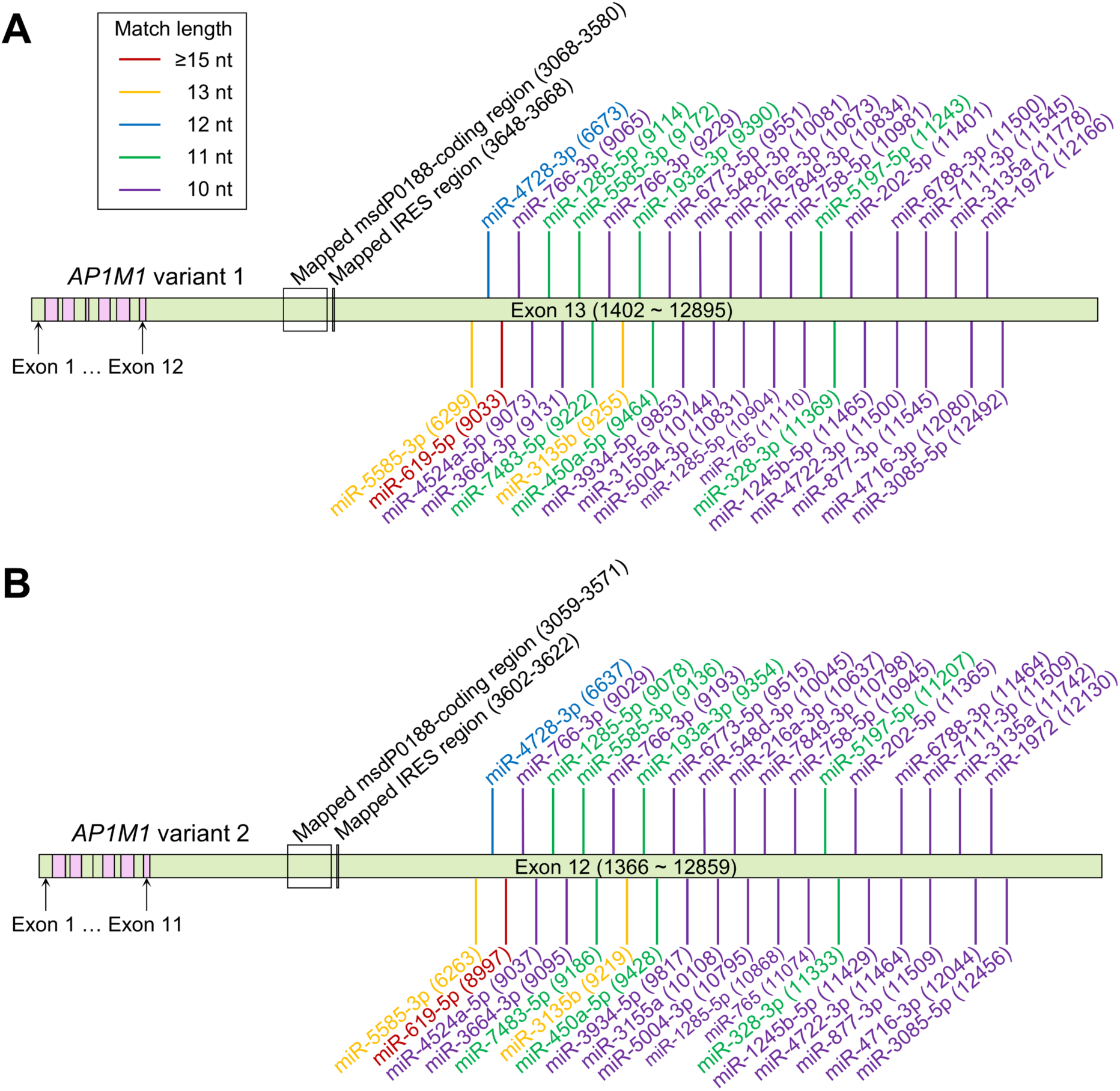
Example msRNAs bound to the *AP1M1* mRNA with 10–15 nt matches. (**A**) In variant 1, all ≥10-nt matches are shown after position 9033, with only the ≥12-nt matches displayed before this position. (**B**) In variant 2, all ≥10-nt matches are shown after position 8997, with only the ≥12-nt matches displayed before this position.

**Fig. S11.**
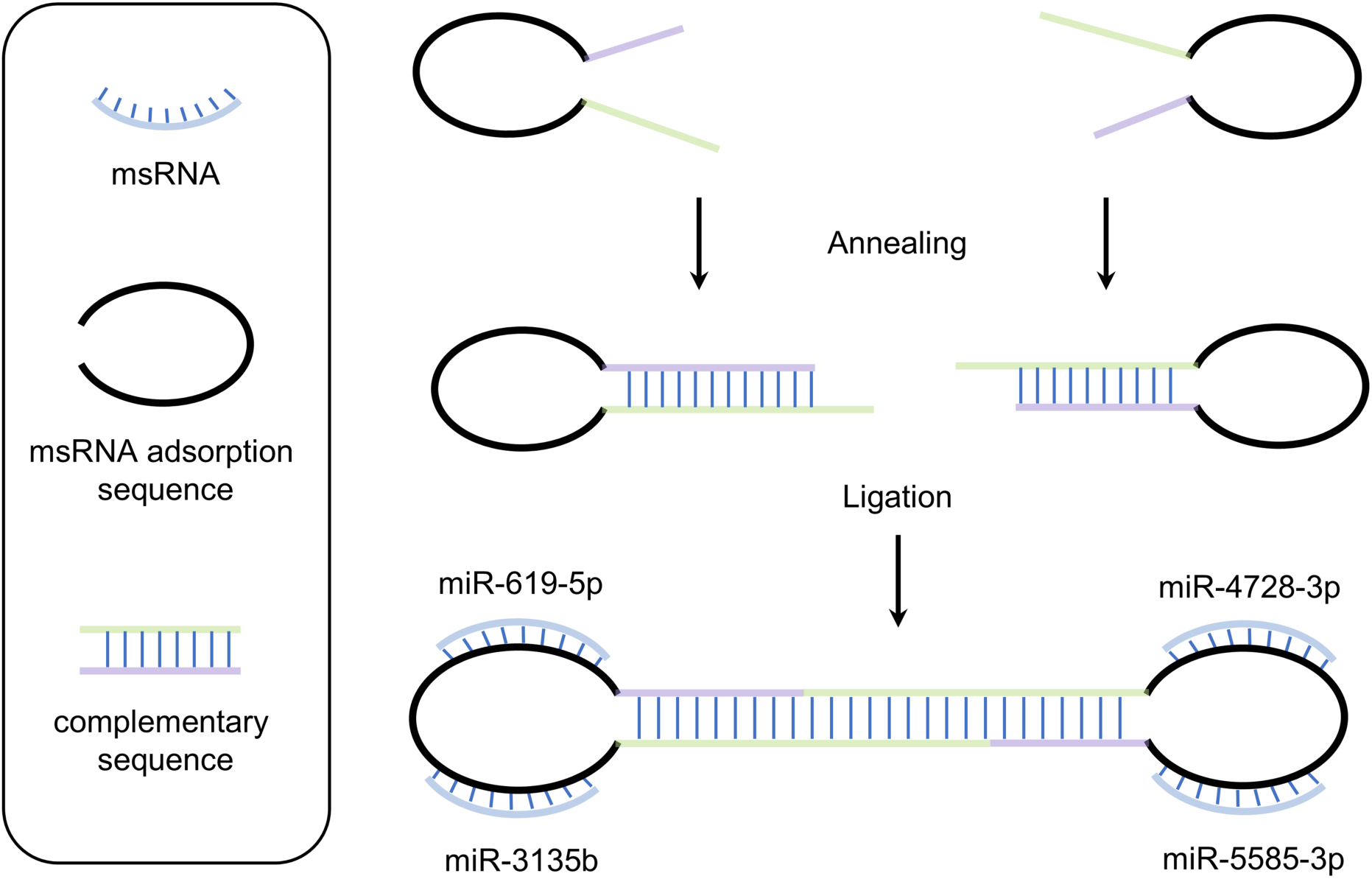
Diagram illustrating the design of CSSD4. CSSD4 is designed to simultaneously bind *miR-619-5p*, *miR-3135b*, *miR-4728-3p*, and *miR-5585-3p*. Both CSSD4 (containing multiple complementary binding sites for msRNAs) and control CSSD (with a scrambled DNA sequence) consist of two DNA half-rings. Single-stranded DNAs were initially synthesized and annealed to form the half-rings, which were then treated with T4 DNA ligase to create fully closed circular DNA structures.

**Fig. S12.**
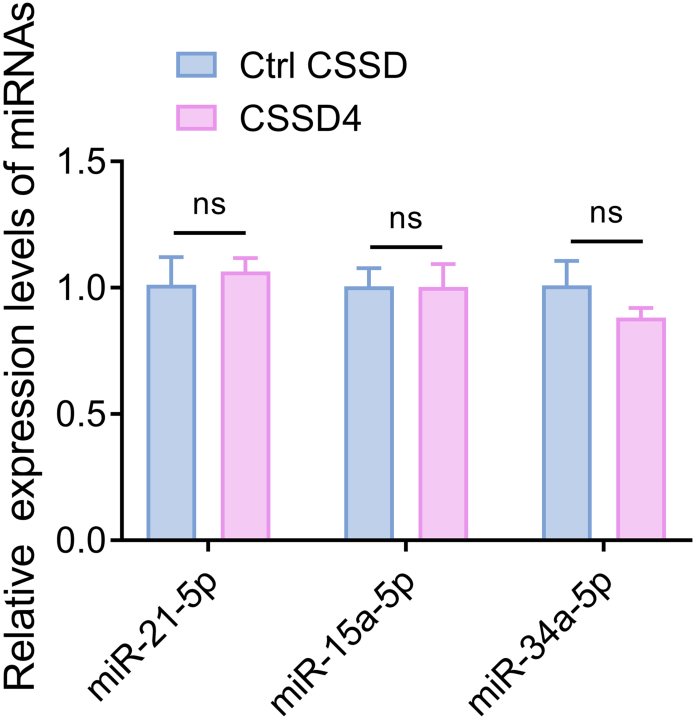
Expression levels of *miR-21-5p*, *miR-15a-3p*, and *miR-34a-5p* in A549 cells as determined by RT-qPCR following CSSD transfection. Error bars represent s.e.m. from n = 3 independent biological replicates. ns, not significant; Student’s *t*-test.

**Fig. S13.**
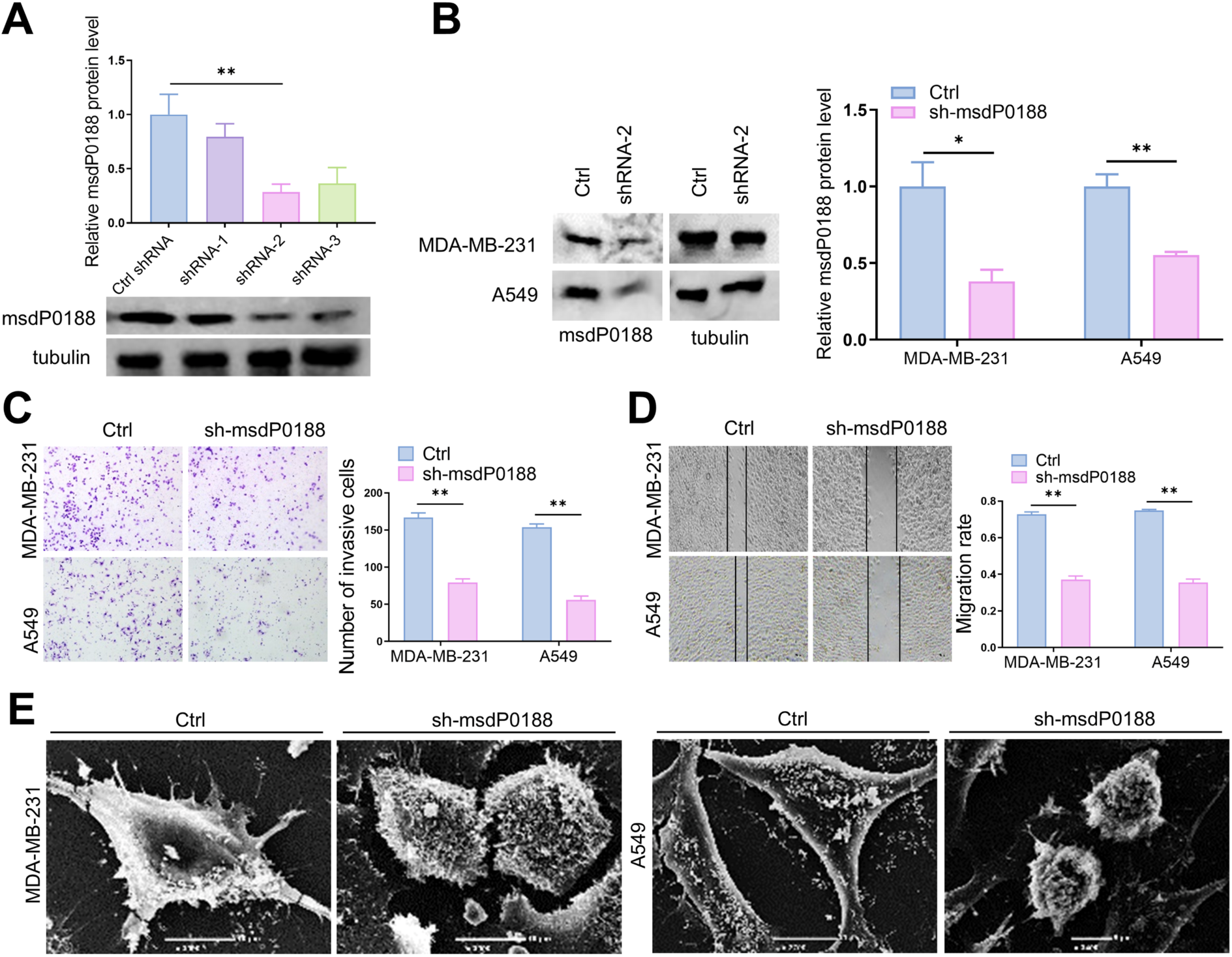
Downregulation of msdP0188 via shRNA transfection inhibits the progression of lung and breast cancer cells in vitro. (**A**) A549 cells were transfected with three interfering lentiviruses targeting msdR0188, along with a control lentivirus carrying a scrambled sequence. The msdP0188 protein level was assessed by Western blot to compare the efficacy of the three shRNAs. (**B**) Downregulation of msdP0188 levels in A549 and MDA-MB-231 cells. (**C**) Effects of sh-msdP0188 on the invasion of A549 and MDA-MB-231 cells. (**D**) Effects of sh-msdP0188 on the migration of A549 and MDA-MB-231 cells. (**E**) Visualization of cell phenotype using scanning electron microscopy following sh-msdP0188 transfection. Error bars represent s.e.m. from n = 3 independent biological replicates. \**P* < 0.05; \*\**P* < 0.01 (Student’s *t*-test).

**Fig. S14.**
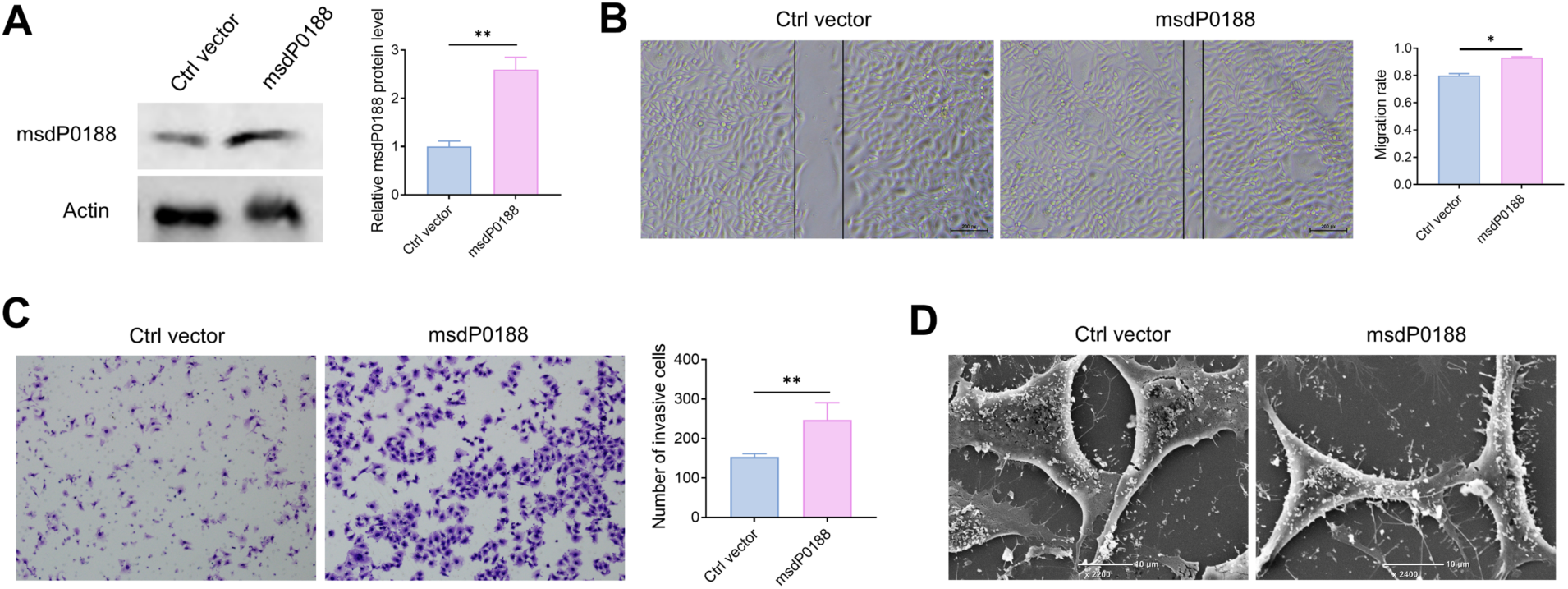
Overexpression of msdP0188 promotes migration and invasion of A549 cells. A549 cells were transfected with either an empty vector (Ctrl) or the PiggyBac Dual promoter plasmid containing the msdP0188 coding sequence. (**A**) Western blot analysis to detect msdP0188 protein levels. (**B**) Effects of msdP0188 overexpression on A549 cell migration. (**C**) Effects of msdP0188 overexpression on A549 cell invasion. (**D**) Scanning electron microscopy analysis of phenotypic changes in A549 cells due to msdP0188 overexpression. Error bars represent s.e.m. from n = 3 independent biological replicates. \**P* < 0.05; \*\**P* < 0.01 (Student’s *t*-test).

**Fig. S15.**
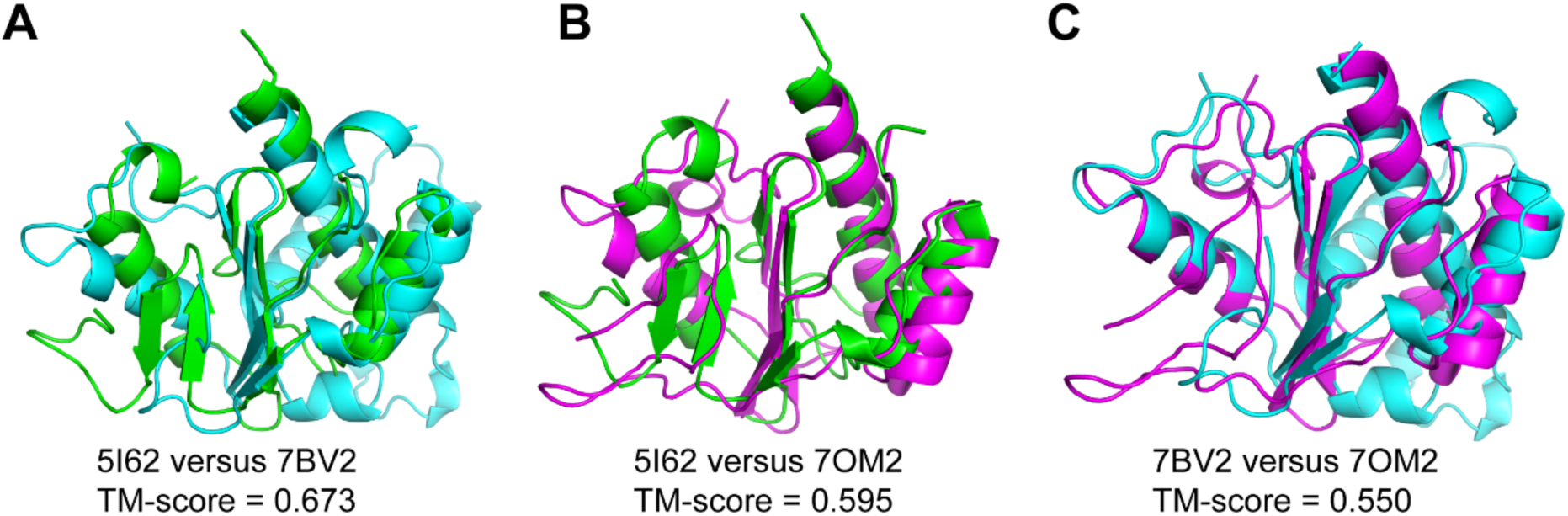
Comparison of the *palm* subdomains of three viral RdRPs. (**A**) Human picobirnavirus strain Hy005102 (green, PDB ID: 5I62) versus SARS-CoV-2 (cyan, PDB ID: 7BV2). (**B**) Human picobirnavirus strain Hy005102 (green) versus *Thosea asigna* virus (magenta, PDB ID: 7OM2). (**C**) SARS-CoV-2 (cyan) versus *Thosea asigna* virus (magenta). The *palm* subdomain of human picobirnavirus strain Hy005102 spans amino acids 231–267 and 325–414. The *palm* subdomain of SARS-CoV-2 spans amino acids 581–627 and 687–812. The *palm* subdomain of *Thosea asigna* virus spans amino acids 304–374 and 444–519.

**Fig. S16.**
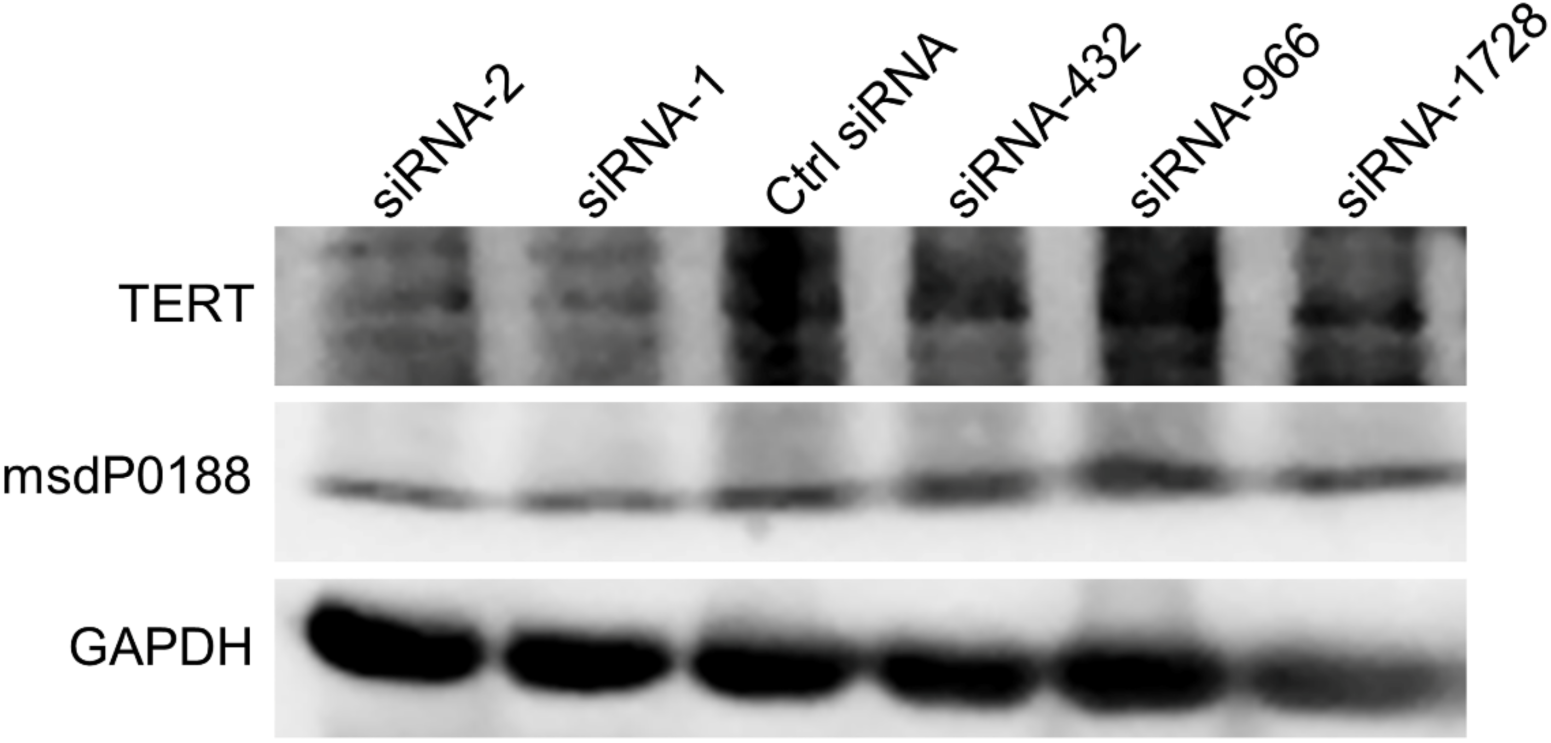
Knockdown of human telomerase reverse transcriptase (TERT) using different siRNAs.

**Fig. S17.**
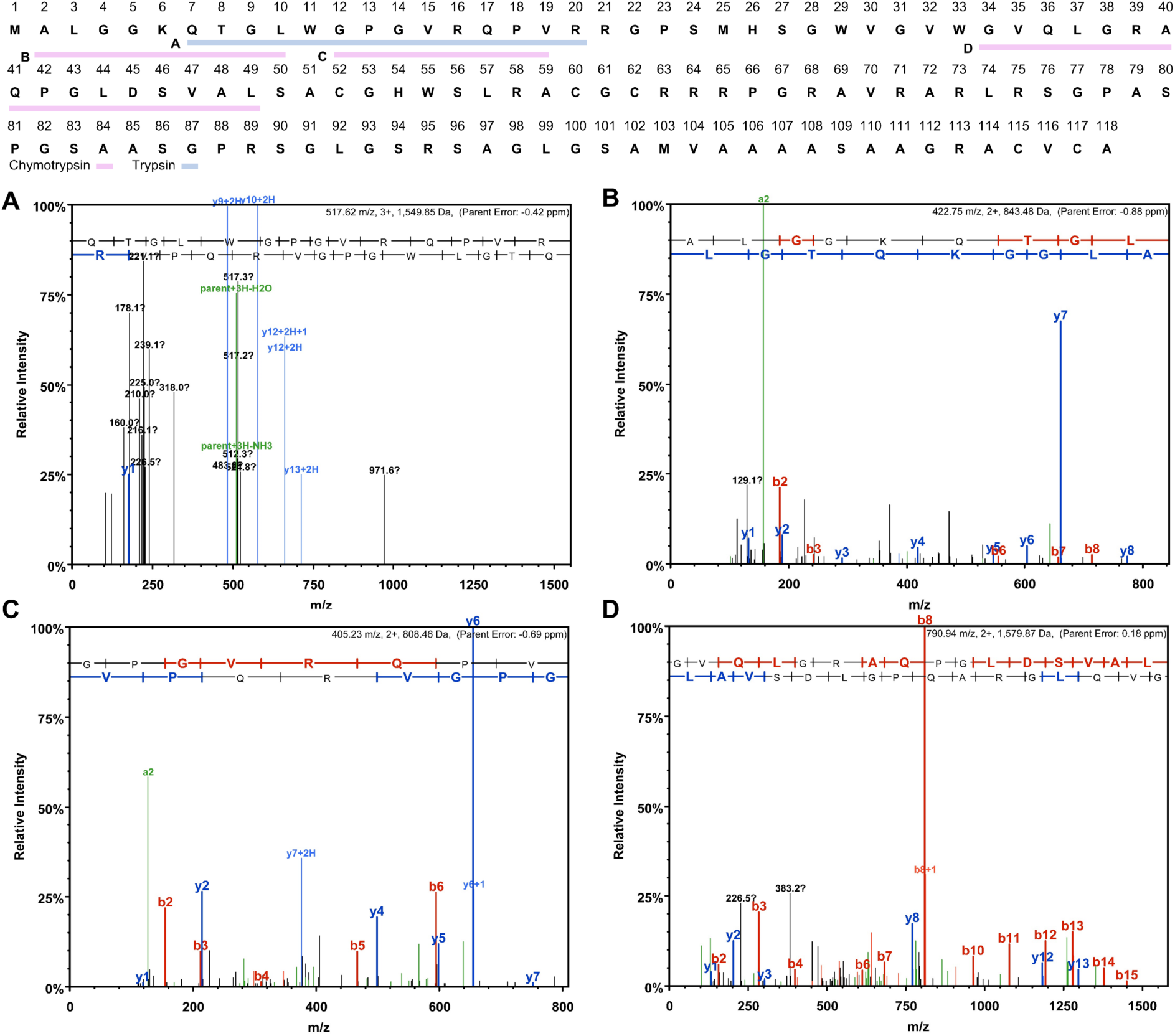
Mass spectrometry assay authenticates msdP0032 peptide fragments after protease digestion. The protein bound to the anti-msdP0032 polyclonal antibody during immunoprecipitation was separated via SDS-PAGE and silver staining to identify a potential target band. This band was excised and subjected to digestion individually with two proteases. The digested samples were then analyzed using mass spectrometry. Four peptide fragments within msdP0032 were validated, including one from trypsin digestion (blue) and three from chymotrypsin digestion (pink).

**Fig. S18.**
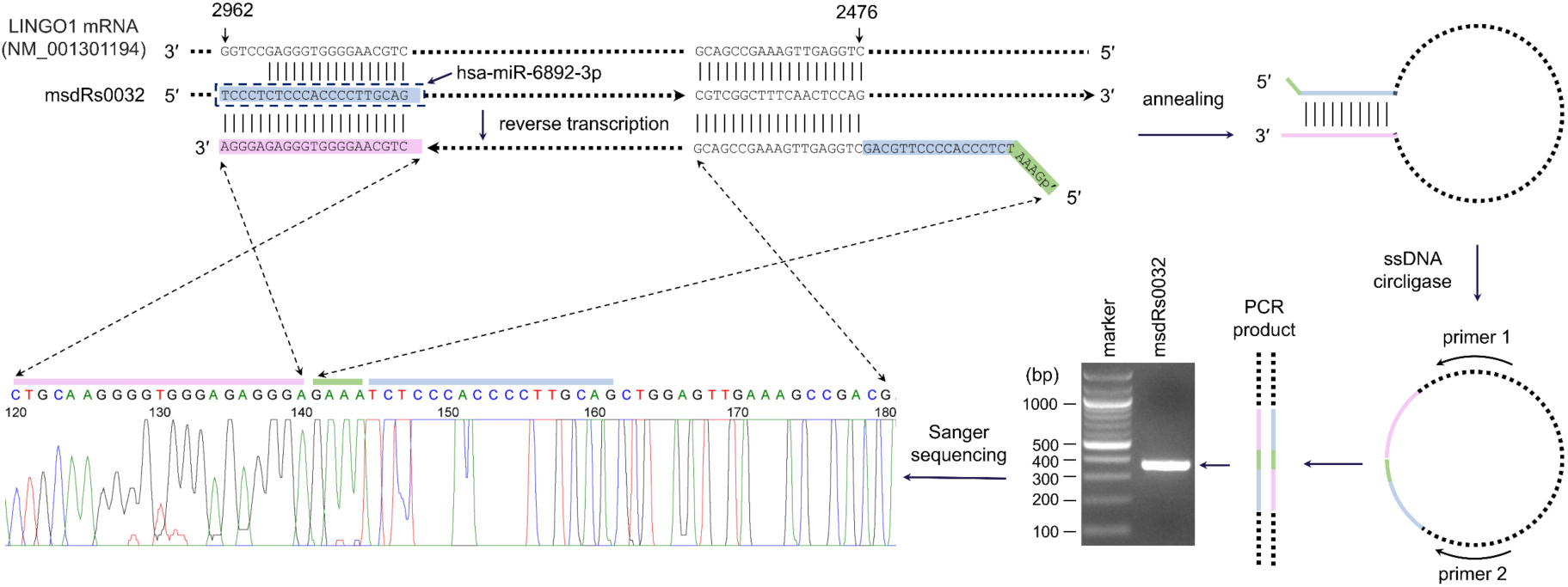
Verification of the 5’-end sequence of msdRs0032 using ssDNA circligase-dependent 5’ RACE.

**Table S1.**
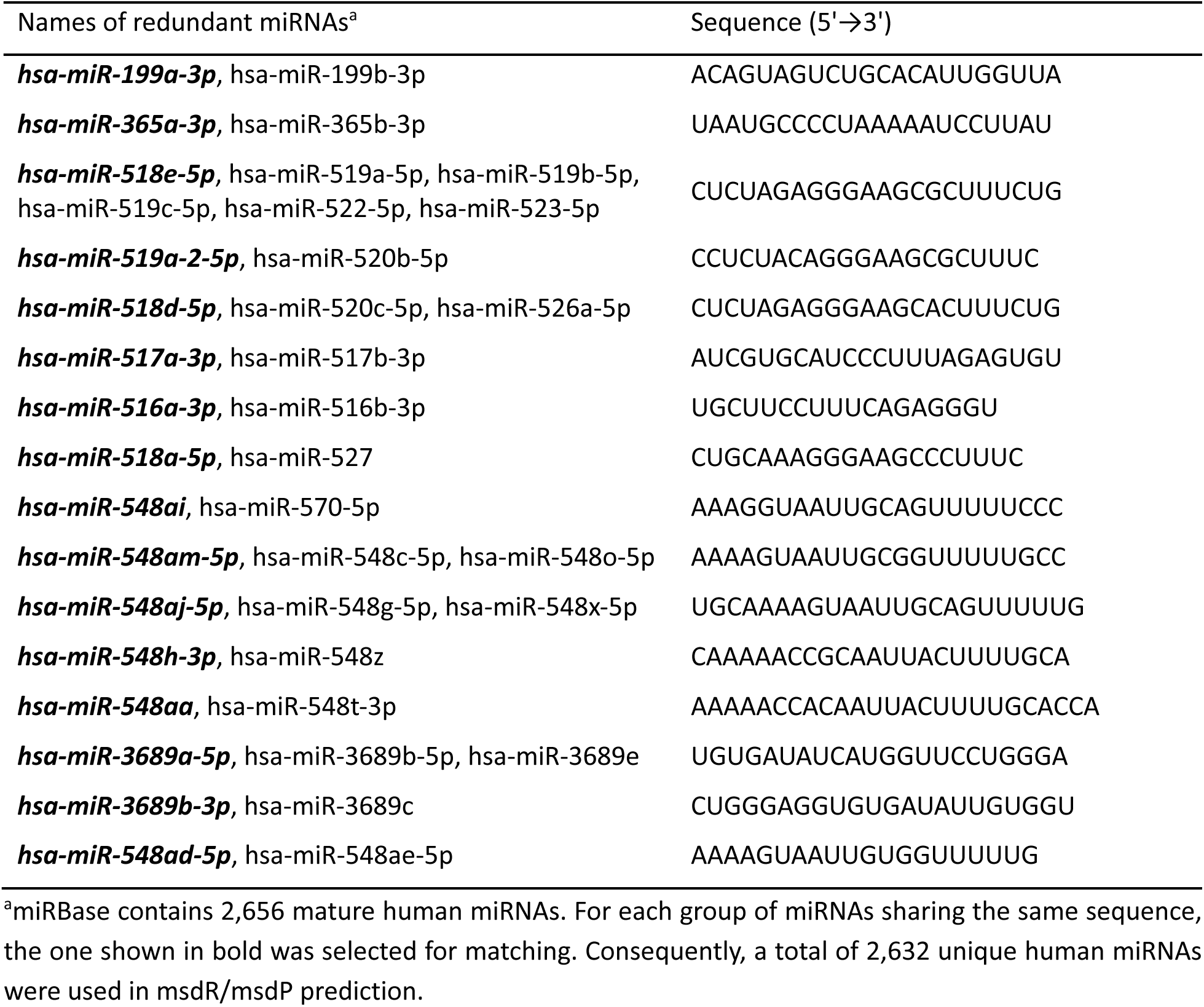
Summary of redundant miRNA sequences in miRBase.

**Table S2.** Summary of number of matched RNAs and predicted msdRs and msdPs.

| Match length (nt) | Number of matched RNAs<sup>a</sup> | Number of unique predicted msdRs<sup>b</sup> | Number of unique (redundant) predicted msdPs<sup>c</sup> |
| --- | --- | --- | --- |
| 15 | 12,953 | 11,121 | 1,239 (5,369) |
| 14 | 28,531 | 24,048 | 2,135 (9,629) |
| 13 | 80,754 | 66,794 | 4,014 (22,829) |
| 12 | 243,976 | 200,386 | 4,252 (34,225) |
| 11 | 828,305 | 679,585 | 4,397 (71,208) |
| 10 | 2,932,303 | 2,399,265 | 3,784 (162,694) |

<sup>a</sup>miRNAs within the NCBI RefSeq RNA database were excluded from the matched RNAs.
<sup>b</sup>Number of redundant msdRs is identical to that of matched RNAs (the 2<sup>nd</sup> column). Unique predicted msdRs stand for non-redundant msdR sequences.
<sup>c</sup>Theoretically, msdPs derived from shorter matches will always include those from longer matches, but this may not be true in practice because NCBI BLAST is a heuristic algorithm which means it may not report all possible alignments, and therefore some hits from longer matches may be missing as the msdR database's size increases.

**Table S3.**
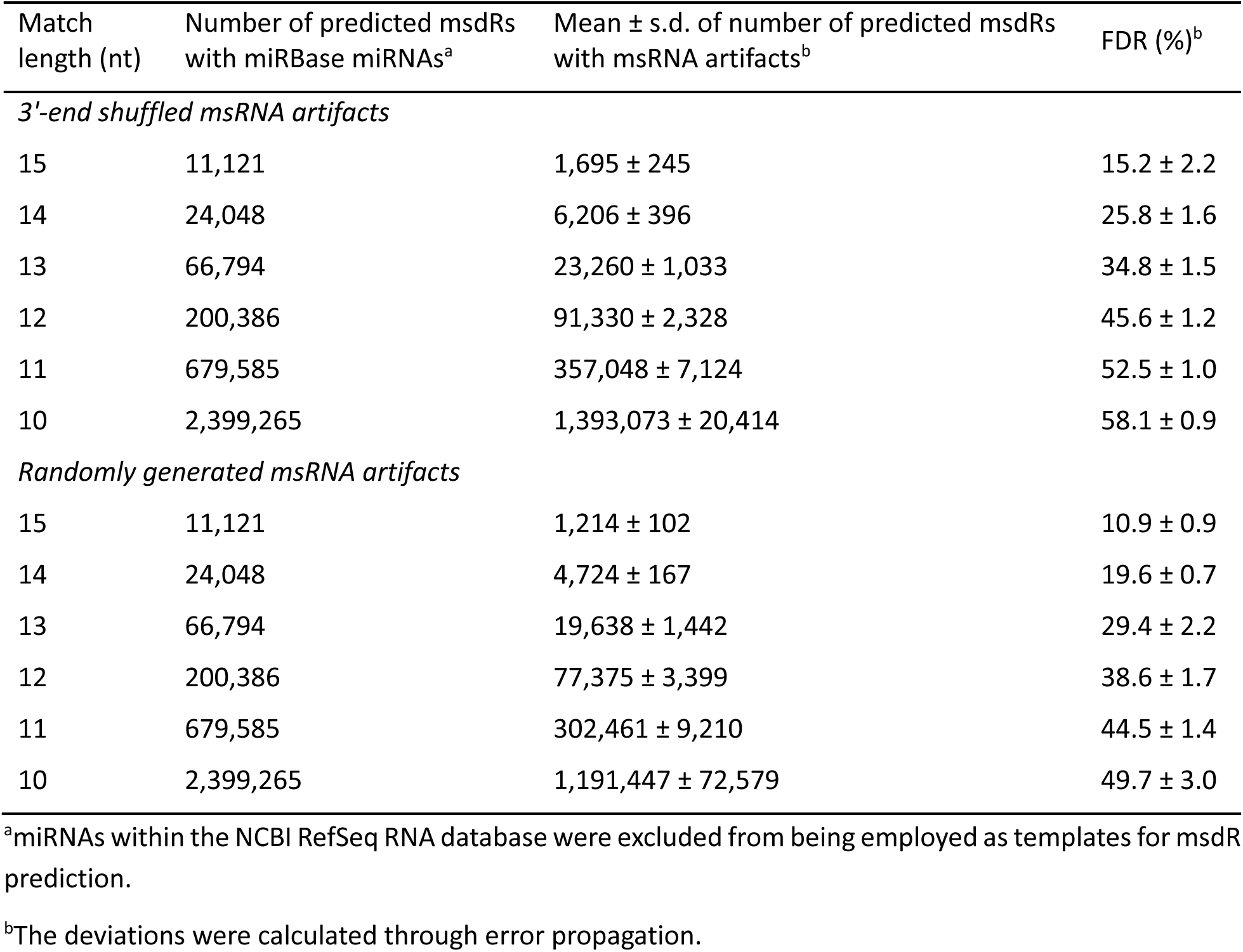
Estimation of potential false discovery rates (FDRs) for msdR prediction using 3’-end shuffled msRNA artifacts based on miRBase miRNAs (n = 2,632) or randomly generated msRNAs (n = 2,632).

**Table S4.**
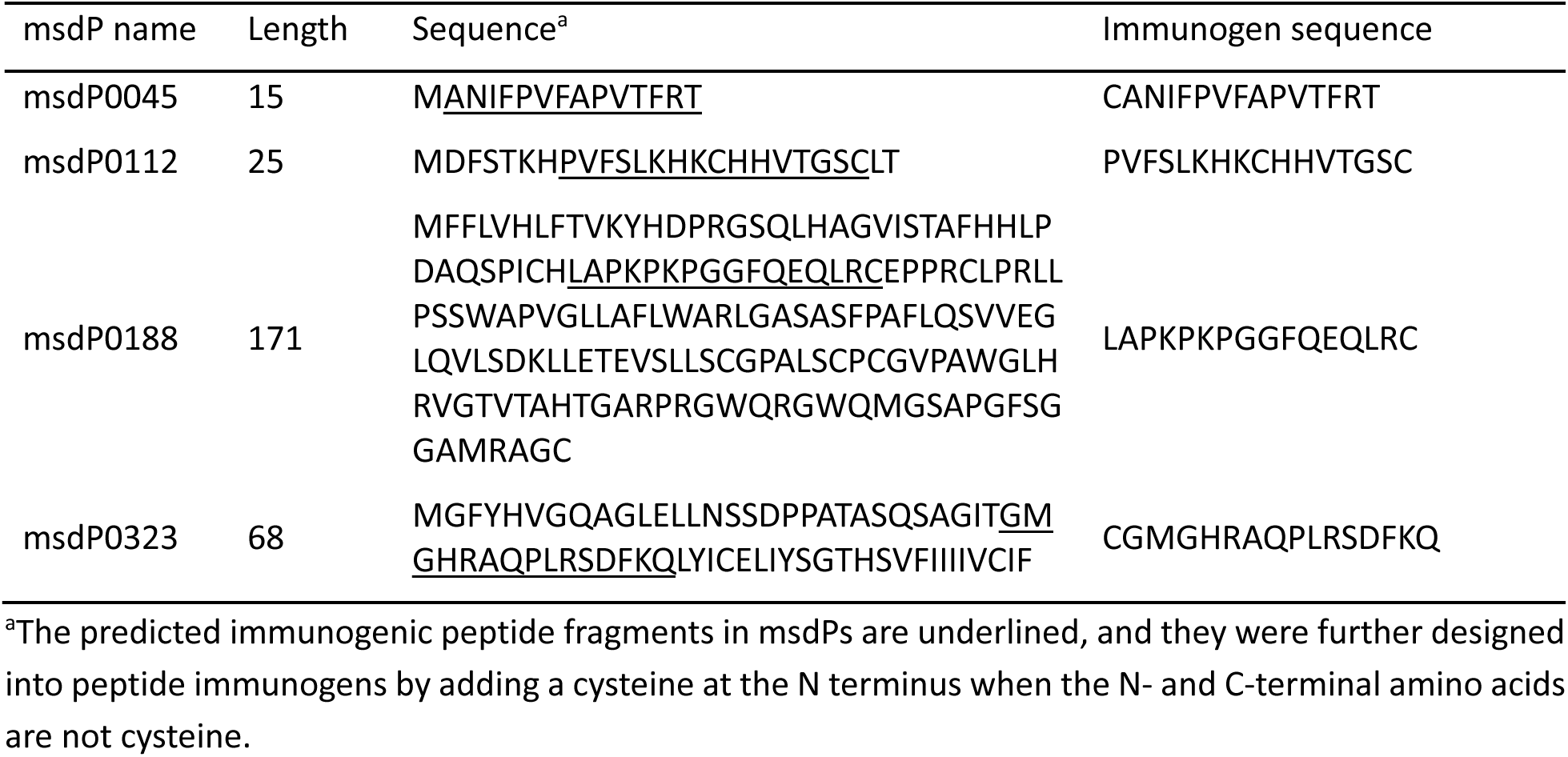
Summary of selected msdPs for immunohistochemistry assay.

**Table S5.** Summary of identifiers and names for selected msdPs and msdRs.

| msdP name | msdP and msdR identifier(s)<sup>a</sup> | msdR name<sup>b</sup> |
| --- | --- | --- |
| msdP0045 | msdP.1370471854(RPS6KB1).414, *msdR.hsa-miR-1972.1370471854(RPS6KB1).2511* | msdRs0045 |
| msdP0112 | msdP.1370451977(NAXE).1372, *msdR.hsa-miR-5585-3p.1370451977(NAXE).3203* | msdRs0112 |
| msdP0188 | msdP.1677479768(AP1M1).3571, *msdR.hsa-miR-619-5p.1677479768(AP1M1).8997*;<br>msdP.1677502062(AP1M1).3607, *msdR.hsa-miR-619-5p.1677502062(AP1M1).9033* | msdRs0188 |
| msdP0323 | msdP.1819229406(C9orf129).1643, *msdR.hsa-miR-3686.1819229406(C9orf129).4888* | msdRs0323 |

<sup>a</sup>The nomenclature of msdP and msdR identifiers can be referred to in Methods. Different msdR sequences may encode the same msdP sequence; such msdRs can be derived from the elongation of the same miRNA bound to mRNA variants, or the elongation of different miRNAs bound to the same RNA. For instance, the *AP1M1* variants 1 and 2 have 12,895 and 12,859 nt, respectively; their only difference is that the 36-nt exon 6 in variant 1 is sliced in variant 2, resulting in a 36-nt shift of the follow-up exons in variant 2 (see **Fig. S10**). For each miRNA able to bind to variant 1 at a position after exon 6, it can also bind to variant 2 at a corresponding position; thus, two *AP1M1* variants can yield similar msdRs which may encode the same protein with different identifiers by our nomenclature (e.g., both msdP.1677479768(AP1M1).3571 and msdP.1677502062(AP1M1).3607 are identifiers for msdP0188).
<sup>b</sup>The number in msdRs name was set identical to that in msdP name for convenience of data presentation.

**Table S6.**
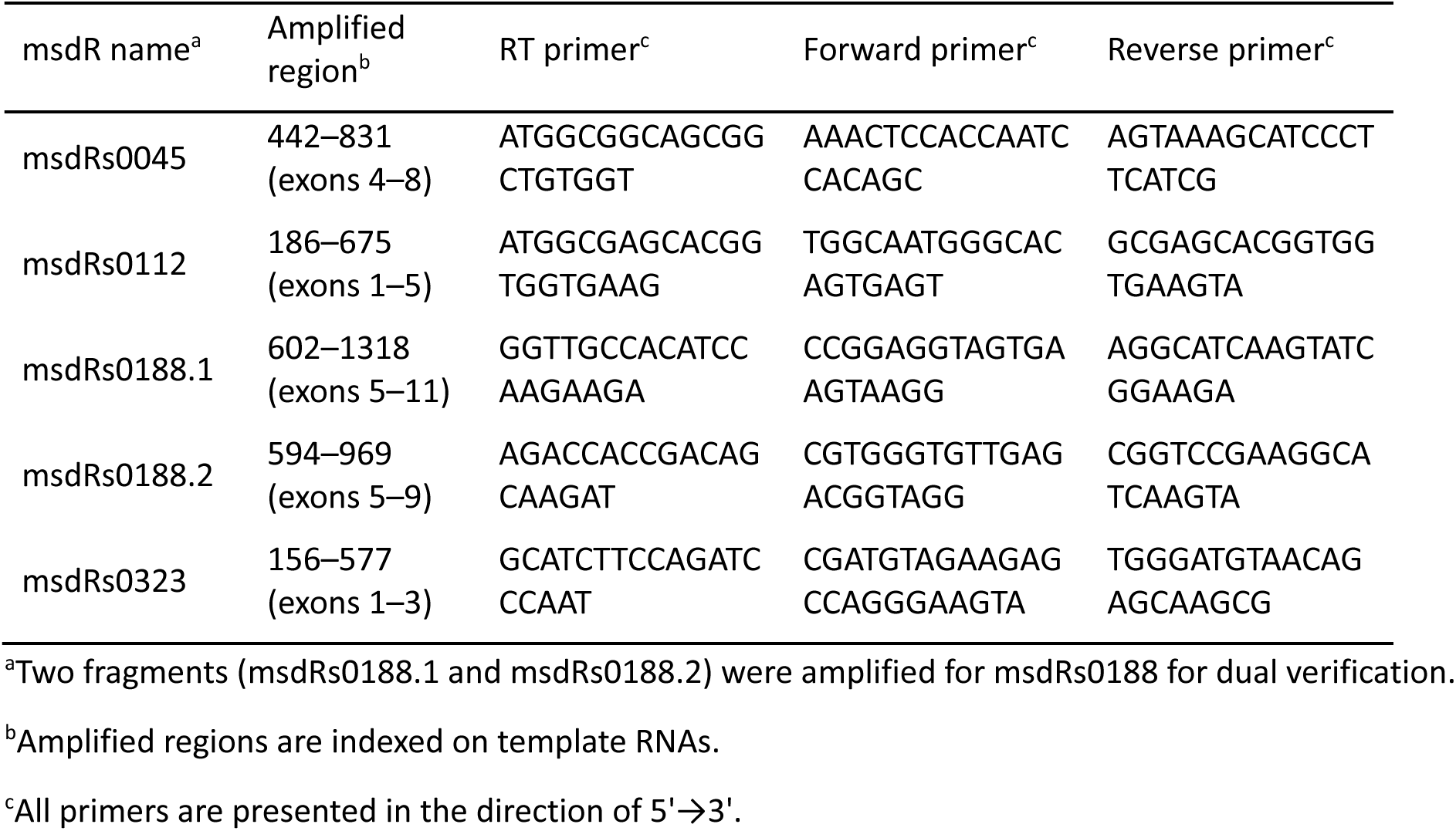
Summary of RT-PCR amplified regions of selected msdRs.

**Table S7.**
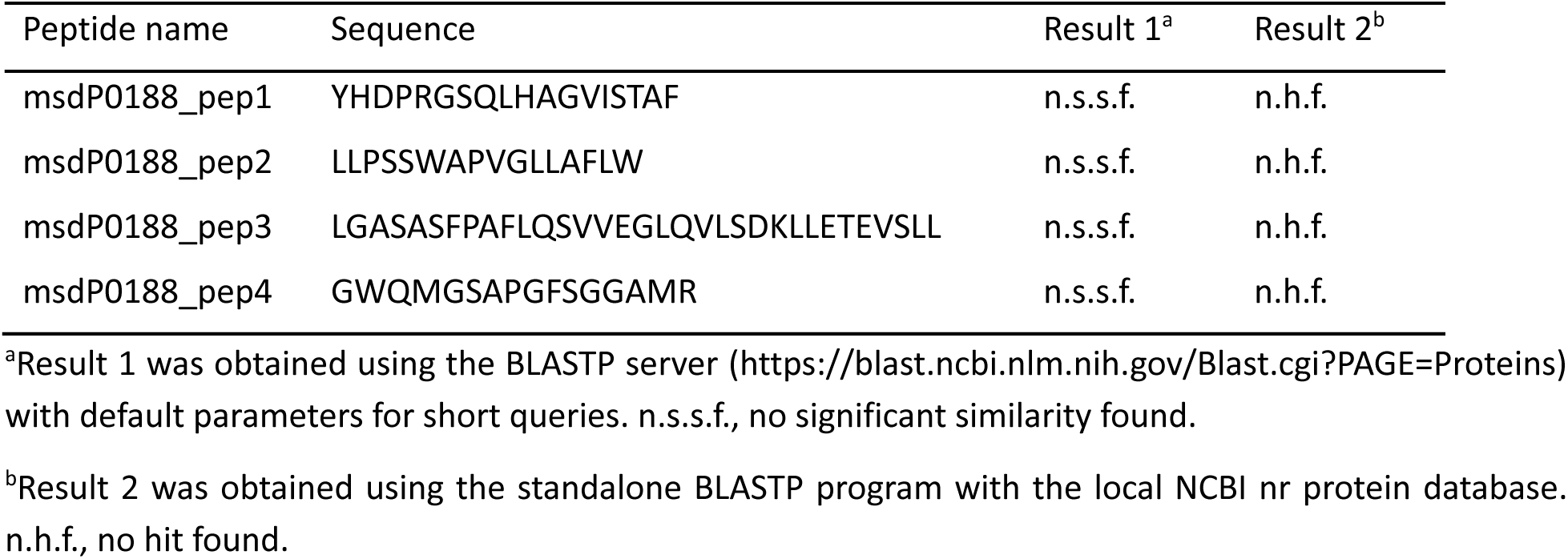
BLASTP search results for four peptides within msdP0188.

**Table S8.**
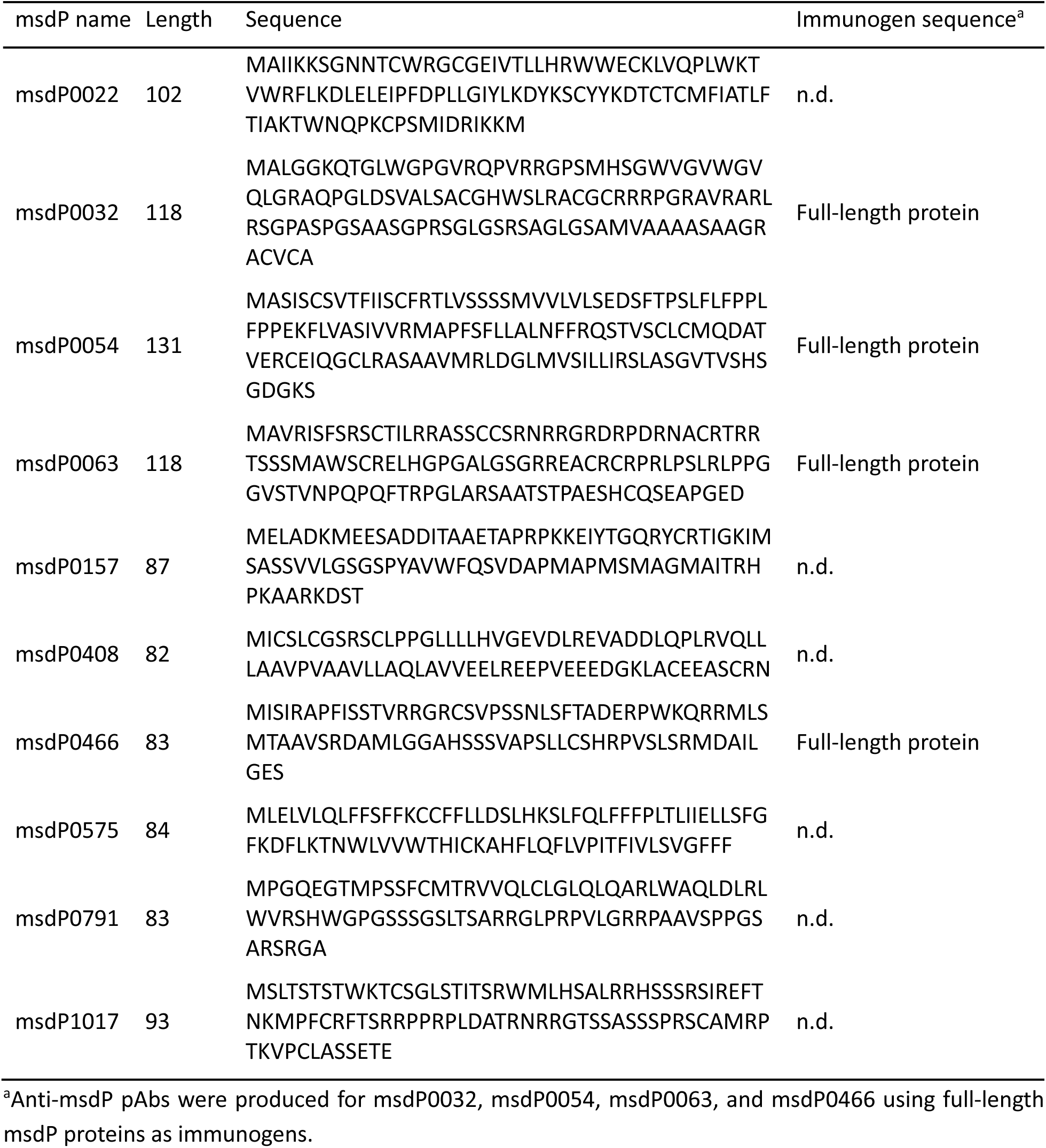
Summary of ten additional msdPs.

**Table S9.** Summary of identifiers and names for ten additional msdPs and their corresponding msdRs.

| msdP name | msdP and msdR identifier(s)<sup>a</sup> | msdR name<sup>b</sup> |
| --- | --- | --- |
| msdP0022 | msdP.1675178226(PCDHA4).4623, *msdR.hsa-miR-619-5p.1675178226(PCDHA4).6660* | msdRs0022 |
| msdP0032 | msdP.1677499153(LINGO1).374, *msdR.hsa-miR-6892-3p.1677499153(LINGO1).2942* | msdRs0032 |
| msdP0054 | msdP.1395168541(CAMK2A).1416, *msdR.hsa-miR-2392.1395168541(CAMK2A).1973*;<br>msdP.1731160493(CAMK2A).1230, *msdR.hsa-miR-2392.1731160493(CAMK2A).1816*;<br>msdP.1889522348(CAMK2A).1230, *msdR.hsa-miR-2392.1889522348(CAMK2A).1787* | msdRs0054 |
| msdP0063 | msdP.1595488117(ALDH3A2).355, *msdR.hsa-miR-619-5p.1595488117(ALDH3A2).2492*;<br>msdP.1676318966(ALDH3A2).355, *msdR.hsa-miR-619-5p.1676318966(ALDH3A2).2617* | msdRs0063 |
| msdP0157 | msdP.1034615633(SLC5A7).2145, *msdR.hsa-miR-8485.1034615633(SLC5A7).4115*;<br>msdP.1034615635(SLC5A7).925, *msdR.hsa-miR-8485.1034615635(SLC5A7).2895*;<br>msdP.1370478493(SLC5A7).4654, *msdR.hsa-miR-8485.1370478493(SLC5A7).6624*;<br>msdP.1653961459(SLC5A7).1364, *msdR.hsa-miR-8485.1653961459(SLC5A7).3334*;<br>msdP.1677499324(SLC5A7).1361, *msdR.hsa-miR-8485.1677499324(SLC5A7).3331*;<br>msdP.1677499965(SLC5A7).1135, *msdR.hsa-miR-8485.1677499965(SLC5A7).3105*;<br>msdP.1677530541(SLC5A7).1327, *msdR.hsa-miR-8485.1677530541(SLC5A7).3297* | msdRs0157 |
| msdP0408 | msdP.228480308(BEND3P3).1375, *msdR.hsa-miR-619-5p.228480308(BEND3P3).2934* | msdRs0408 |
| msdP0466 | msdP.1523709028(NELFA).251, *msdR.hsa-miR-2276-3p.1523709028(NELFA).1190* | msdRs0466 |
| msdP0575 | msdP.1034655404(SEPTIN14).1241, *msdR.hsa-miR-619-5p.1034655404(SEPTIN14).1559*;<br>msdP.1519473604(SEPTIN14).1220, *msdR.hsa-miR-619-5p.1519473604(SEPTIN14).1538* | msdRs0575 |
| msdP0791 | msdP.1519312642(DCHS1).408, *msdR.hsa-miR-6752-5p.1519312642(DCHS1).7852* | msdRs0791 |
| msdP1017 | msdP.1675178226(PCDHA4).497, *msdR.hsa-miR-619-5p.1675178226(PCDHA4).6660* | msdRs1017 |

**Table S10.**
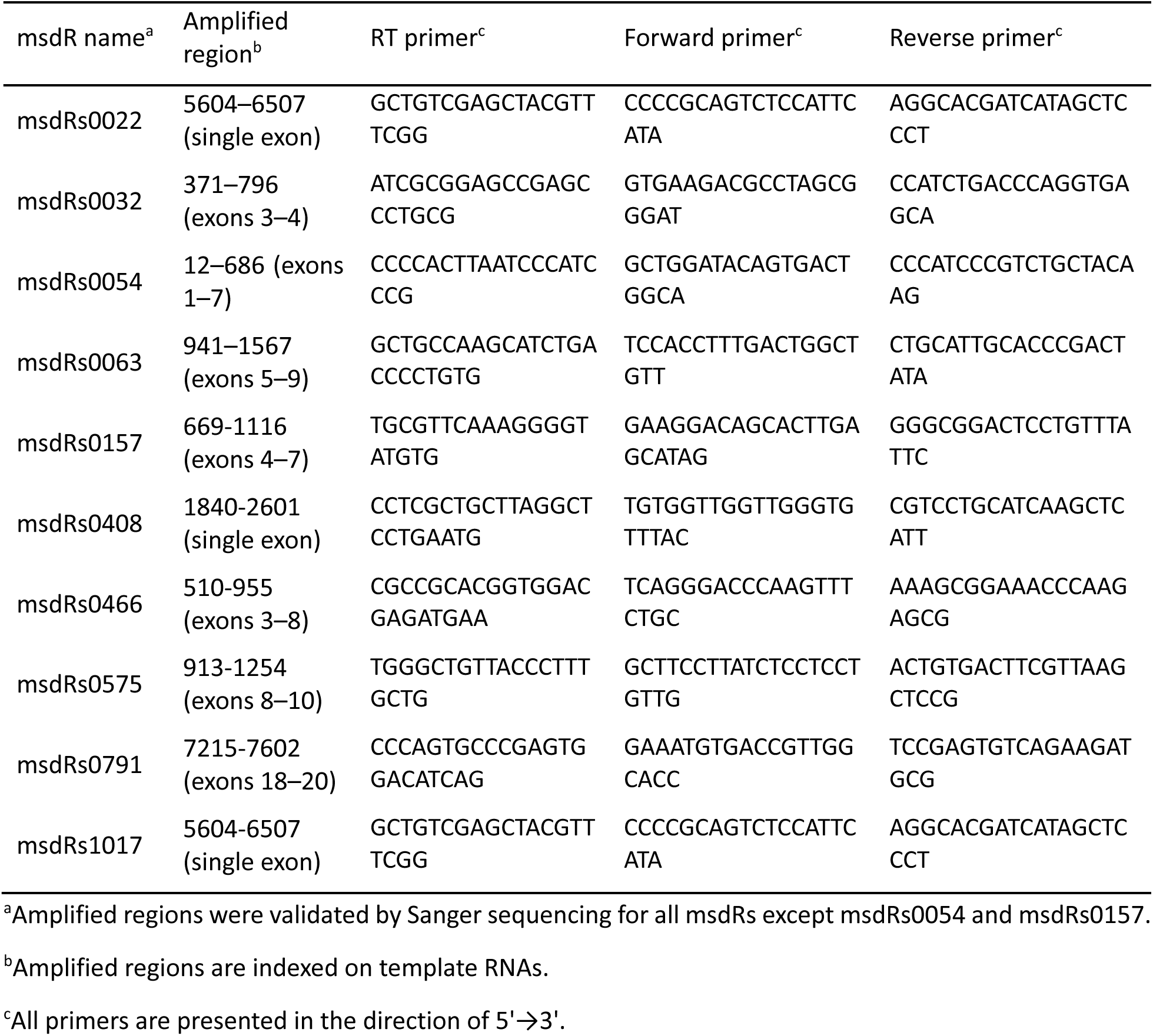
Summary of RT-PCR amplified regions of ten additional msdRs.

**Table S11.**
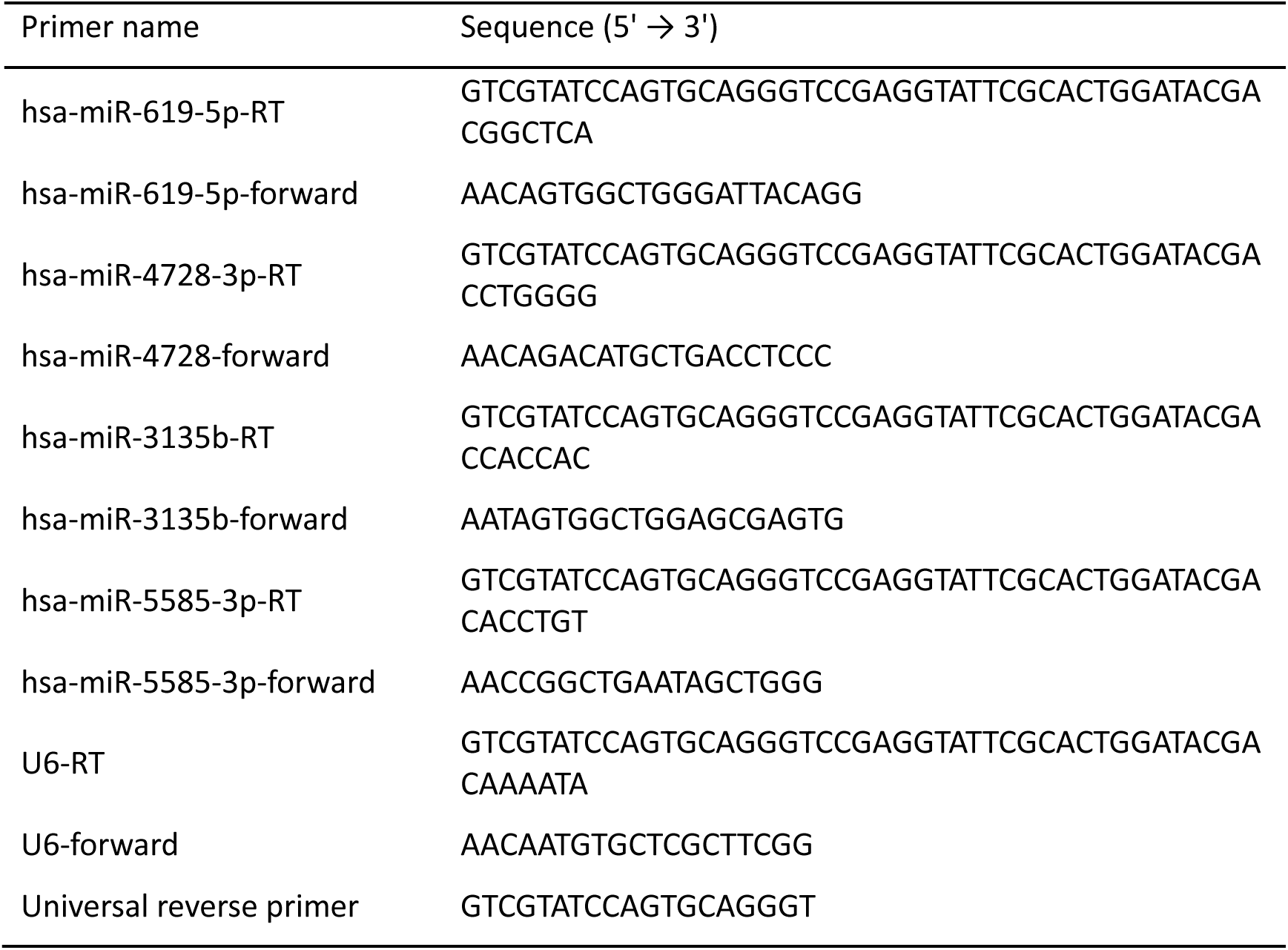
Summary of RT-qPCR primers for msRNA qualification.

## Other Supplementary Data

**Data S1**. Summary of unique miRBase human miRNAs (n = 2,632) used in this study.

**Data S2**. Unique msdRs predicted using 10–15 nt matches between msRNA and template RNAs.

**Data S3**. Unique msdPs translated from msdRs.

**Data S4**. Simulated msRNA artifacts for estimating the false discovery rates for msdR prediction.

**Data S5**. Summary of the number of msdRs predicted using either database-documented miRNAs or control msRNA artifacts.

**Data S6**. Summary of 46 msdPs with homologs in the NCBI nr protein database.

**Data S7**. Summary of six msdPs with identical hit sequences in the NCBI nr protein database.

**Data S8**. Summary of 1,193 msdPs.

**Data S9**. Summary of 941 msdPs matching with mass spectra from cancer cell lines.

**Data S10**. Summary of 555 msdPs with confident peptide-spectrum matches via PepQuery2 search.

**Data S11**. Sanger sequencing data.

**Data S12**. Mass spectrometry data.

**Data S13**. Summary of 358 D*x*DGD motif-containing human proteins with more than 500 amino acids.

**Data S14**. Summary of 33 human proteins in the PDB with TM-score greater than 0.5 to the *palm* subdomain of three viral RdRPs.

**Data S15**. Summary of 84 human proteins in the AlphaFold Protein Structure Database with TM-score greater than 0.5 to the *palm* subdomain of three viral RdRPs.

Data S1 to S15 are publicly available at https://doi.org/10.5281/zenodo.7826531.

## References

1 Lee, R. C., Feinbaum, R. L. & Ambros, V. The C. elegans heterochronic gene lin-4 encodes small RNAs with antisense complementarity to lin-14. Cell 75, 843–854 (1993). 10.1016/0092-8674(93)90529-y

2 Wightman, B., Ha, I. & Ruvkun, G. Posttranscriptional regulation of the heterochronic gene lin-14 by lin-4 mediates temporal pattern formation in C. elegans. Cell 75, 855–862 (1993). 10.1016/0092-8674(93)90530-4

3 Lu, T. X. & Rothenberg, M. E. MicroRNA. J Allergy Clin Immunol 141, 1202–1207 (2018). 10.1016/j.jaci.2017.08.034

4 Brennan, G. P. & Henshall, D. C. MicroRNAs as regulators of brain function and targets for treatment of epilepsy. Nat Rev Neurol 16, 506–519 (2020). 10.1038/s41582-020-0369-8

5 Kim, T. & Croce, C. M. MicroRNA and ER stress in cancer. Semin Cancer Biol 75, 3–14 (2021). 10.1016/j.semcancer.2020.12.025

6 Lee, T. J. et al. Strategies to Modulate MicroRNA Functions for the Treatment of Cancer or Organ Injury. Pharmacol Rev 72, 639–667 (2020). 10.1124/pr.119.019026

7 Tay, Y., Zhang, J., Thomson, A. M., Lim, B. & Rigoutsos, I. MicroRNAs to Nanog, Oct4 and Sox2 coding regions modulate embryonic stem cell differentiation. Nature 455, 1124–1128 (2008). 10.1038/nature07299

8 Vasudevan, S., Tong, Y. & Steitz, J. A. Switching from repression to activation: microRNAs can up-regulate translation. Science 318, 1931–1934 (2007). 10.1126/science.1149460

9 Orom, U. A., Nielsen, F. C. & Lund, A. H. MicroRNA-10a binds the 5’UTR of ribosomal protein mRNAs and enhances their translation. Mol Cell 30, 460–471 (2008). 10.1016/j.molcel.2008.05.001

10 Krischuns, T., Lukarska, M., Naffakh, N. & Cusack, S. Influenza Virus RNA-Dependent RNA Polymerase and the Host Transcriptional Apparatus. Annu Rev Biochem 90, 321–348 (2021). 10.1146/annurev-biochem-072820-100645

11 Gao, Y. et al. Structure of the RNA-dependent RNA polymerase from COVID-19 virus. Science 368, 779–782 (2020). 10.1126/science.abb7498

12 O’Leary, N. A. et al. Reference sequence (RefSeq) database at NCBI: current status, taxonomic expansion, and functional annotation. Nucleic Acids Res. 44, D733–745 (2016). 10.1093/nar/gkv1189

13 Altschul, S. F. et al. Gapped BLAST and PSI-BLAST: a new generation of protein database search programs. Nucleic Acids Res. 25, 3389–3402 (1997).

14 Zhao, J. et al. IRESbase: A Comprehensive Database of Experimentally Validated Internal Ribosome Entry Sites. Genomics Proteomics Bioinformatics 18, 129–139 (2020). 10.1016/j.gpb.2020.03.001

15 Kozomara, A., Birgaoanu, M. & Griffiths-Jones, S. miRBase: from microRNA sequences to function. Nucleic Acids Res. 47, D155–D162 (2019). 10.1093/nar/gky1141

16 Lewis, B. P., Shih, I. H., Jones-Rhoades, M. W., Bartel, D. P. & Burge, C. B. Prediction of mammalian microRNA targets. Cell 115, 787–798 (2003). 10.1016/s0092-8674(03)01018-3

17 Brennecke, J., Stark, A., Russell, R. B. & Cohen, S. M. Principles of microRNA-target recognition. PLoS Biol. 3, e85 (2005). 10.1371/journal.pbio.0030085

18 Eng, J. K., McCormack, A. L. & Yates, J. R. An approach to correlate tandem mass spectral data of peptides with amino acid sequences in a protein database. J Am Soc Mass Spectrom 5, 976–989 (1994). 10.1016/1044-0305(94)80016-2

19 Wen, B. & Zhang, B. PepQuery2 democratizes public MS proteomics data for rapid peptide searching. Nat Commun 14, 2213 (2023). 10.1038/s41467-023-37462-4

20 Meng, J. et al. Derepression of co-silenced tumor suppressor genes by nanoparticle-loaded circular ssDNA reduces tumor malignancy. Sci Transl Med 10 (2018). 10.1126/scitranslmed.aao6321

21 Wu, H. et al. G-quadruplex-enhanced circular single-stranded DNA (G4-CSSD) adsorption of miRNA to inhibit colon cancer progression. Cancer Med 12, 9774–9787 (2023). 10.1002/cam4.5721

22 Iyer, L. M., Koonin, E. V. & Aravind, L. Evolutionary connection between the catalytic subunits of DNA-dependent RNA polymerases and eukaryotic RNA-dependent RNA polymerases and the origin of RNA polymerases. BMC Struct. Biol. 3, 1 (2003). 10.1186/1472-6807-3-1

23 Zhang, Y. & Skolnick, J. TM-align: a protein structure alignment algorithm based on the TM-score. Nucleic Acids Res. 33, 2302–2309 (2005).

24 Berman, H. M. et al. The Protein Data Bank. Acta Crystallographica Section D Biological Crystallography 58, 899–907 (2002). 10.1107/s0907444902003451

25 Jumper, J. et al. Highly accurate protein structure prediction with AlphaFold. Nature 596, 583–589 (2021). 10.1038/s41586-021-03819-2

26 Tunyasuvunakool, K. et al. Highly accurate protein structure prediction for the human proteome. Nature 596, 590–596 (2021). 10.1038/s41586-021-03828-1

27 Maida, Y. et al. An RNA-dependent RNA polymerase formed by TERT and the RMRP RNA. Nature 461, 230–235 (2009). 10.1038/nature08283

28 Maida, Y., Yasukawa, M. & Masutomi, K. De Novo RNA Synthesis by RNA-Dependent RNA Polymerase Activity of Telomerase Reverse Transcriptase. Mol Cell Biol 36, 1248–1259 (2016). 10.1128/MCB.01021-15

29 Takahashi, M. et al. Eribulin penetrates brain tumor tissue and prolongs survival of mice harboring intracerebral glioblastoma xenografts. Cancer Sci. 110, 2247–2257 (2019). 10.1111/cas.14067

30 Machitani, M. et al. Maintenance of R-loop structures by phosphorylated hTERT preserves genome integrity. Nat. Cell Biol. 26, 932–945 (2024). 10.1038/s41556-024-01427-6

31 Zhang, X., Mar, V., Zhou, W., Harrington, L. & Robinson, M. O. Telomere shortening and apoptosis in telomerase-inhibited human tumor cells. Genes Dev 13, 2388–2399 (1999). 10.1101/gad.13.18.2388

32 Mocellin, S. et al. Telomerase reverse transcriptase locus polymorphisms and cancer risk: a field synopsis and meta-analysis. J. Natl. Cancer Inst. 104, 840–854 (2012). 10.1093/jnci/djs222

33 Kapranov, P. et al. New class of gene-termini-associated human RNAs suggests a novel RNA copying mechanism. Nature 466, 642–646 (2010). 10.1038/nature09190

34 Shi, K., Liu, T., Fu, H., Li, W. & Zheng, X. Genome-wide analysis of lncRNA stability in human. PLoS Comput Biol 17, e1008918 (2021). 10.1371/journal.pcbi.1008918

35 Clark, M. B. et al. Genome-wide analysis of long noncoding RNA stability. Genome Res 22, 885–898 (2012). 10.1101/gr.131037.111

36 Swartwout, S. G., Preisler, H., Guan, W. D. & Kinniburgh, A. J. Relatively stable population of c-myc RNA that lacks long poly(A). Mol Cell Biol 7, 2052–2058 (1987). 10.1128/mcb.7.6.2052-2058.1987

37 Kim, Y. et al. PKR Senses Nuclear and Mitochondrial Signals by Interacting with Endogenous Double-Stranded RNAs. Mol Cell 71, 1051–1063 e1056 (2018). 10.1016/j.molcel.2018.07.029

38 Kim, Y. et al. PKR is activated by cellular dsRNAs during mitosis and acts as a mitotic regulator. Genes Dev 28, 1310–1322 (2014). 10.1101/gad.242644.114

39 Gregersen, L. H. et al. MOV10 Is a 5’ to 3’ RNA helicase contributing to UPF1 mRNA target degradation by translocation along 3’ UTRs. Mol Cell 54, 573–585 (2014). 10.1016/j.molcel.2014.03.017

40 Koh, H. R., Xing, L., Kleiman, L. & Myong, S. Repetitive RNA unwinding by RNA helicase A facilitates RNA annealing. Nucleic Acids Res 42, 8556–8564 (2014). 10.1093/nar/gku523

41 Sharma, D., Putnam, A. A. & Jankowsky, E. Biochemical Differences and Similarities between the DEAD-Box Helicase Orthologs DDX3X and Ded1p. J Mol Biol 429, 3730–3742 (2017). 10.1016/j.jmb.2017.10.008

42 Song, H. & Ji, X. The mechanism of RNA duplex recognition and unwinding by DEAD-box helicase DDX3X. Nat Commun 10, 3085 (2019). 10.1038/s41467-019-11083-2

43 Omenn, G. S. et al. The 2024 Report on the Human Proteome from the HUPO Human Proteome Project. J. Proteome Res. 23, 5296–5311 (2024). 10.1021/acs.jproteome.4c00776

44 Okamoto, N. et al. Maintenance of tumor initiating cells of defined genetic composition by nucleostemin. Proc. Natl. Acad. Sci. U.S.A. 108, 20388–20393 (2011).

